# Functional role of respiratory supercomplexes in mice: segmentation of the Q_pool_ and SCAF1

**DOI:** 10.1101/826115

**Authors:** Enrique Calvo, Sara Cogliati, Pablo Hernansanz-Agustín, Marta Loureiro-López, Adela Guarás, Rafael A. Casuso, Fernando García-Marqués, Rebeca Acín-Pérez, Yolanda Martí-Mateos, JC. Silla-Castro, Marta Carro-Alvarellos, Jesús R. Huertas, Jesús Vázquez, J.A. Enríquez

## Abstract

Mitochondrial respiratory complexes assemble into different forms of supercomplexes (SC). In particular, SC III_2_+IV require the SCAF1 protein. However, the structural role of this factor in the formation of the respirasome (I+III_2_+IV) and the physiological role of SCs are controversial. Here, we study C57BL/6J mice harbouring either non-functional SCAF1, the full knock-out for SCAF1 or the wild-type version of the protein and found a growth and exercise phenotype due to the lack of functional SCAF1. By combining quantitative data-independent proteomics, high resolution 2D Blue Native Gel Electrophoresis and functional analysis of enriched respirasome fractions, we show that SCAF1 confers structural attachment between III_2_ and IV within the respirasome, increases NADH-dependent respiration and reduces ROS production. Furthermore, through the expression of AOX in cells and mice we confirm that CI-CIII superassembly segments the CoQ in two pools and modulates CI-NADH oxidative capacity. These data demonstrate that SC assembly, regulated by SCAF1, modulates the functionality of the electron transport chain.

## Introduction

The mitochondrial cristae are the main site of biological energy conversion through the respiratory complexes I to V known as oxidative phosphorylation system (OXPHOS). Our understanding of the structure of the mitochondria electron transport chain was shaken in 2000 by Herman Schägger when proposing that respiratory complexes could form superstructures called supercomplexes (SCs), among which the ones containing CI, CIII and CIV were named respirasomes (Schagger, 2000). This proposal came from a novel electrophoretic methodology mastered by that author named Blue Native Gel Electrophoresis (BNGE). First received with extreme skepticism, SCs are nowadays generally accepted as true biological entities. They are present in mitochondria from very different sources (Eubel et al., 2003; Schägger, 2002); they are able to respire (Acín-Pérez et al., 2008); specific factors for the formation of some SCs have been discovered (Lapuente-Brun et al., 2013); and the cryo-electron microscopy structures of the respirasome (I+III_2_+IV) and the supercomplex I+III_2_ have been obtained (Gu et al., 2016; Letts et al., 2016; Sousa et al., 2016). Although all these data demonstrate their existence, the physiological role of SCs is still under strong debate. A line of thinking proposes that SCs have no functional role (Milenkovic et al., 2017). Other authors indicate that SCs optimize electron flux to gain efficiency in energy generation while minimizing reactive oxygen species production (Lapuente-Brun et al., 2013; Lenaz and Genova, 2007; Maranzana et al., 2013). We proposed the “Plasticity Model”, where individual and super-assembled complexes coexist in a regulated equilibrium within the inner mitochondrial membrane (Acín-Pérez et al., 2008; Enríquez, 2016). Thus, SCs rearrange in response to a shift in the metabolic source of electrons (Guarás et al., 2016), in the metabolic adaptation of specialised cells in vivo (Garaude et al., 2016), or to adapt the cell to different nutrients and physiological conditions (Greggio et al., 2017; Lopez-Fabuel et al., 2016). For instance, under starvation the preferential use of fatty acids reduces the levels of SCs containing CI, so that free III_2_ is more accessible to electrons coming from FADH_2_ (Lapuente-Brun et al., 2013).

The controversy was centered in the biological role of SCAF1 in respirasomes and in the role of SCs in the functional segmentation of the CoQ pool. In mouse, SCAF1 is 113 amino acid long and has a high homology in its carboxy terminus with one subunit of CIV, for which there are two isoforms (COX7A1-80 aa and COX7A2-83 aa). The amino-terminal portion of the protein has no homology with any known protein. SCAF1 is required to super-assemble CIII and CIV by replacing COX7A2 in the structure of CIV and by binding CIII through its amino-terminal portion (Cogliati et al., 2016). Interestingly, all inbred mouse C57BL/6 sub-strains investigated to date (Enríquez, 2019), harbor a non-functional version of SCAF1 (named SCAF1^111^) that, due to a microdeletion that eliminate two amino acids, is unable to interact with CIV (Cogliati et al., 2016). In the absence of functional SCAF1, the SC III_2_+IV and the majority of SCs containing CI, CIII and CIV cannot be formed (Cogliati et al., 2016; Lapuente-Brun et al., 2013; Lobo-Jarne et al., 2018). However, in some instances, particularly in heart mitochondria from BL/6 mice, a comigration between CI, CIII and CIV suggestive of the presence of respirasomes can be observed (Cogliati et al., 2016; Lobo-Jarne et al., 2018). This raised doubts about the relevance of SCAF1 for the formation and function of the respirasome (Lobo-Jarne et al., 2018). In addition, we observed, confirming the previous suggestions (Bianchi et al., 2004), that superassembly partially segments CoQ in two pools, one predominantly dedicated to FAD and another to NAD(Lapuente-Brun et al., 2013). Noteworthy, these data were obtained using isolated mitochondria. Elsewhere, but using sub-mitochondrial particles, opposite conclusions were reported (Fedor and Hirst, 2018), calling the partial CoQ partitioning hypothesis into question.

In this work we demonstrate that the ablation of SCAF1 decreases the performance of mice under rigorous metabolic stresses such as reduced food supply or intense exercise demand. In addition, by combining data-independent proteomics, 2D BNGE-based structural analysis and functional studies of the BNGE enriched respirasome fractions, we demonstrate that SCAF1 plays a key role in the regulation of the structure of respirasome and its bioenergetics performance. In addition, through the expression of AOX in several cell systems and animals, we have been able to confirm the segmentation of the CoQ in two pools is a consequence of the superassembly between CI and CIII. These findings demonstrate that SCs play a physiological role and provide a molecular mechanism for the phenotype observed in animals.

## Results

### The ablation of functional SCAF1 compromises growth under food restriction and impairs exercise performance

One of the more recurrent arguments to challenge the proposal that SCAF1 and the superassembly between complexes III and IV are bioenergetically relevant is the belief that all C57BL/6 mice sub-strains, which harbor a non-functional SCAF1, lack an apparent phenotype. Since this belief has never been experimentally addressed, we first investigated it in detail. We compared the phenotype of C57BL/6 mice with full ablation of SCAF1 (SCAF1^-/-^ or SCAF1^KO^), with the non-functional version of SCAF1 (SCAF1^111/111^), and with the wild-type and functional version of this gene in homozygosis (SCAF1^113/113^). We first confirmed that SCAF1^113/113^ liver and heart mitochondria express high levels of SCAF1 that are associated substantially with the respirasome and SC: III_2_+IV (Fig. 1A). On the other hand, and as described before (Cogliati et al., 2016), SCAF1^111^ was very unstable and could only be found in minor amounts, interacting with CIII either in the I+III_2_ or in III_2_ (Fig. 1A). In the full KO mice, no SCAF1 could be detected (Fig. 1A). Work from our group revealed that the ablation of SCAF1 in zebrafish affects the growth of male and female animals, the female fertility and the swimming performance of otherwise healthy animals. We also showed that all these phenotypic features disappear when overfeeding the animals (García-Poyatos et al., 2019). Therefore, we hypothesized that since in the usual food regime of mouse facilities animals are fed ad-libitum, a situation that can be considered overfeeding, C57BL/6 mice would be protected against similar manifestations of the lack of functional SCAF1. Therefore, we first evaluated if the presence of functional SCAF1 in mice had any physiological impact under restricted food administration. Animals were feed ad libitum for 24 hours every three days and were fasted during the rest of the time (Fig. 1B). During the first days the animals suffered a period of adaptation in which SCAF1^113^ males were able to maintain or even increase their weight, whereas SCAF1^111^ and KO males showed the opposite trend (Fig. 1C). This period also resulted in the loss of some mice, without significant differences between strains but with a clearly better survival rate of females vs. males (Fig. S1A). Once adapted, all animals started gaining weight. These results indicate that the absence of SCAF^113^ affects mice physiology and that SCAF^111^ does not retain any significant functionality (Fig. 1C). Interestingly, females were insensitive to the presence of functional SCAF1 (Fig. 1C), probably due to a different female mechanism of adaptation to fasting (Della Torre et al., 2018). Next, we evaluated the impact of functional SCAF1 on exercise performance. Thus, we found that both male and female SCAF1^113^ were able to reach a 30% higher maximum speed in the treadmill than any other group of animals (Fig. 1D). These studies demonstrate that the presence of functional SCAF1 has a direct impact on the mice phenotype, implying that supercomplex formation allows a more efficient electron transport chain. This increased efficiency allows males to better extract energy from aliments, and both sexes to respond to higher muscle work demands. The observation that SCAF1 ablation causes such evident phenotype demands a deeper look into the molecular role of this protein in the function of the mitochondrial electron transport chain, which could help to solve the controversy on this issue. To that aim, we combined unbiased complexome analysis by mass spectrometry, 2D-BNGE evaluation of the interaction of respiratory complexes within supercomplexes, and functional analysis of enriched fractions of complexes and supercomplexes in the presence or in the absence of functional SCAF1.

**Figure 1.**
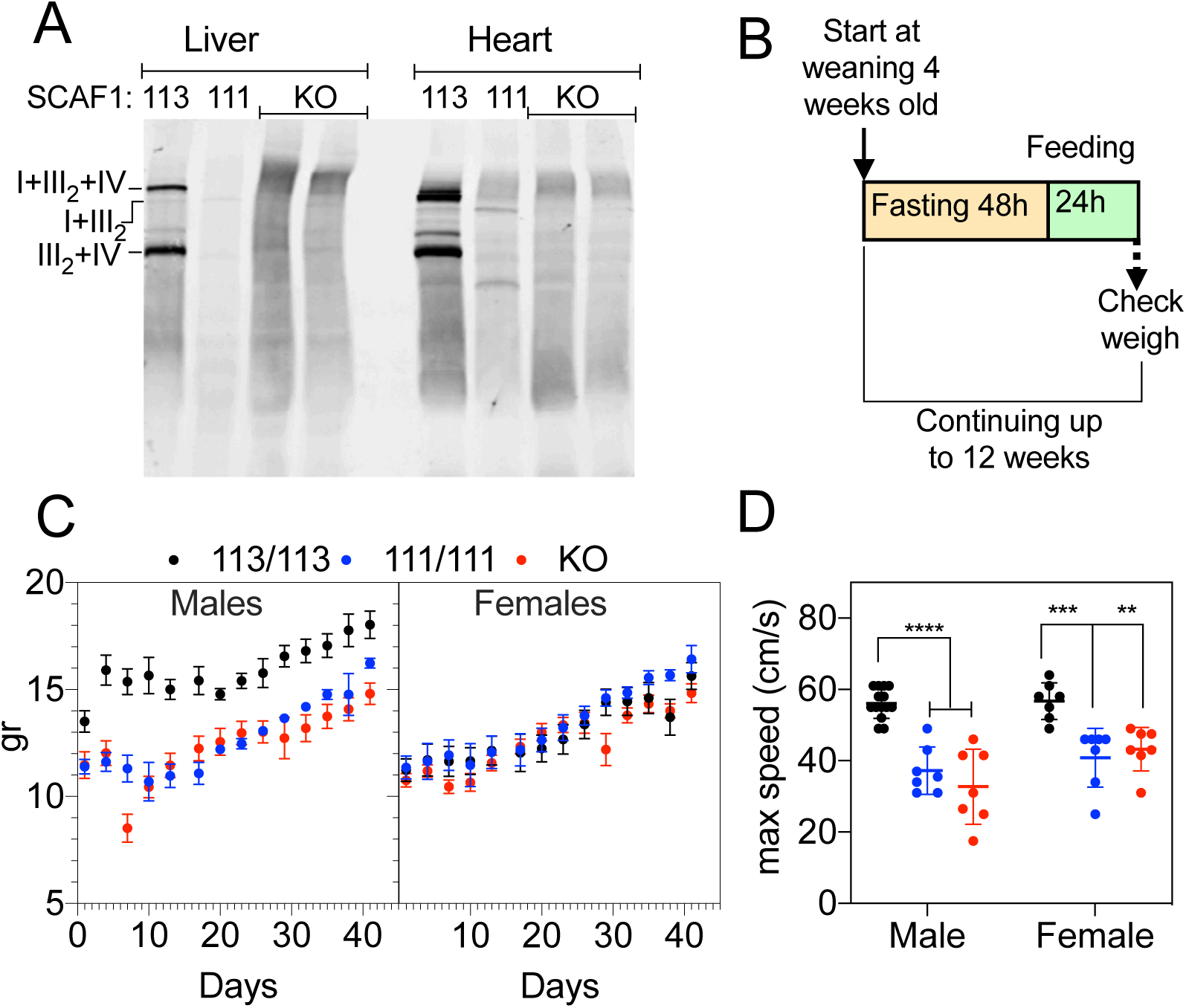
SCAF1 deficiency induces a conspicuous phenotype in mice. A, BNGE followed by western-blot showing the absence or presence of SCAF1 in the indicated tissue and mouse strain. 113: C57BL/6JOlaHsd mice with the functional version of SCAF1; 111: C57BL/6JOlaHsd mice (it harbors a non-functional version of SCAF1); KO C57BL/6JOlaHsd mice without SCAF1. **B,** Scheme of the food restriction experiment analyzed in **C**. **D,** Effect of SCAF1 on the maximum speed running in a treadmill by the indicated mouse groups.

### The size distribution of OXPHOS components is tissue-specific as a consequence of superassembly

First, we applied Blue-DiS, a recently developed technology that takes advantage of the reproducibility and sensitivity of data-independent mass spectrometry(Guarás et al., 2016), to perform mitochondrial complexome profiling from a completely hypothesis-free perspective. Thus, mitochondria-enriched preparations from heart, brain and liver from CD1 mice and from the cell line L929 repopulated with mtDNA from C57BL/6mice (L929^C57^), having all of them the wild type functional version of SCAF1, were separated by BNGE and each lane was cut into 26 slices; each slice was subjected to trypsin protein digestion and analyzed by MS using the DiS method we previously developed (Guarás et al., 2016). We identified 1,134 proteins classified as mitochondrial in the Mitocarta 2.0 (Calvo et al., 2016), which correspond to 98% of all Mitocarta annotated proteins (Fig. S1B). Mitochondrial enrichment was different between samples, being heart (81% of peptide identifications annotated in Mitocarta) and liver (71%) more enriched in known mitochondrial proteins; while brain showed a lower enrichment (38%) (Fig. S1C). In all cases an almost complete coverage of components of OXPHOS complexes was achieved (Dataset 1). Every subunit from CII and CIII was identified (Fig. S1D), together with 41 out of the 44 annotated proteins from CI (Fig. S1D). Concerning CIV, this complex is built out of 14 proteins but several of them have isoforms making up a total of 21 possible components. However, COX6B2 and COX7B2 are both expressed only in sperm, and COX4i2 only in lung, so that 18 proteins were potentially identifiable in our samples, from which we detected 16 (Fig. S1D). Regarding CV, we could detect 17 out of the 20 structural proteins of this complex (Fig. S1D). Apparent molecular weights could be accurately assigned to each one of the 26 BNGE slices from the known masses of individual complexes and of SC (Fig. S1E).

We performed a cross-correlation analysis of protein abundances across different slices within the same mitochondrial type (Fig. S2A). This analysis showed that the relative abundances of all proteins belonging to either CI, CIII, CIV or CV were constant across slices (Fig. S2A & Supplementary Table 1), and that the most representative slice from each complex also had the same protein proportions across the different mitochondrial types (Fig. S2B & Supplementary Table 2). In striking contrast, the electrophoretic mobility protein distributions of CI, CIII and CIV, but not of CV, were markedly different from one mitochondrial source to another (Fig. S2C, D). Thus, complexes in different tissues have the same protein composition but form tissue-specific high molecular weight structures.

Notably, the larger structures (slides 1-8) consistently contained identical proportions of CI and CIII, or CIII and CIV proteins (Fig. 2A, B). Cross correlation analysis confirmed that the protein composition in the I+III+IV, I+III and III+IV structures was maintained between the adjacent slices (Fig. 2C) and between tissues (Fig. 2D), reflecting the formation of SC in an unbiased manner. To obtain a quantitative estimate of the distribution of these structures in each tissue we modelled the complete migration profiles of CI, CIII and CIV by unsupervised gaussian deconvolution. We found that in all mitochondrial types the profile of CI and CIII could be accurately explained as a superimposition of broad gaussian peaks corresponding to a ternary complex (I+III+IV), to binary I+III and III+IV structures and to free CI, CIII and CIV forms (dashed red lines in Fig. 2E and Fig. S2D), having these components the same electrophoretic mobilities and similar peak widths across all mitochondrial sources (Fig. 2F). We also found that the profile of CIV could not be adequately explained by these structures around slices 6-12 (Fig. 2E, black arrows), suggesting the presence of significant amounts of CIV dimers and multimers. Thus, the addition of two more CIV structures (IV_2_ and IV_m_) was sufficient to model the migration profile of this complex in all mitochondrial types (dashed red lines in Fig. 2E and Fig. S2D).

**Figure 2.**
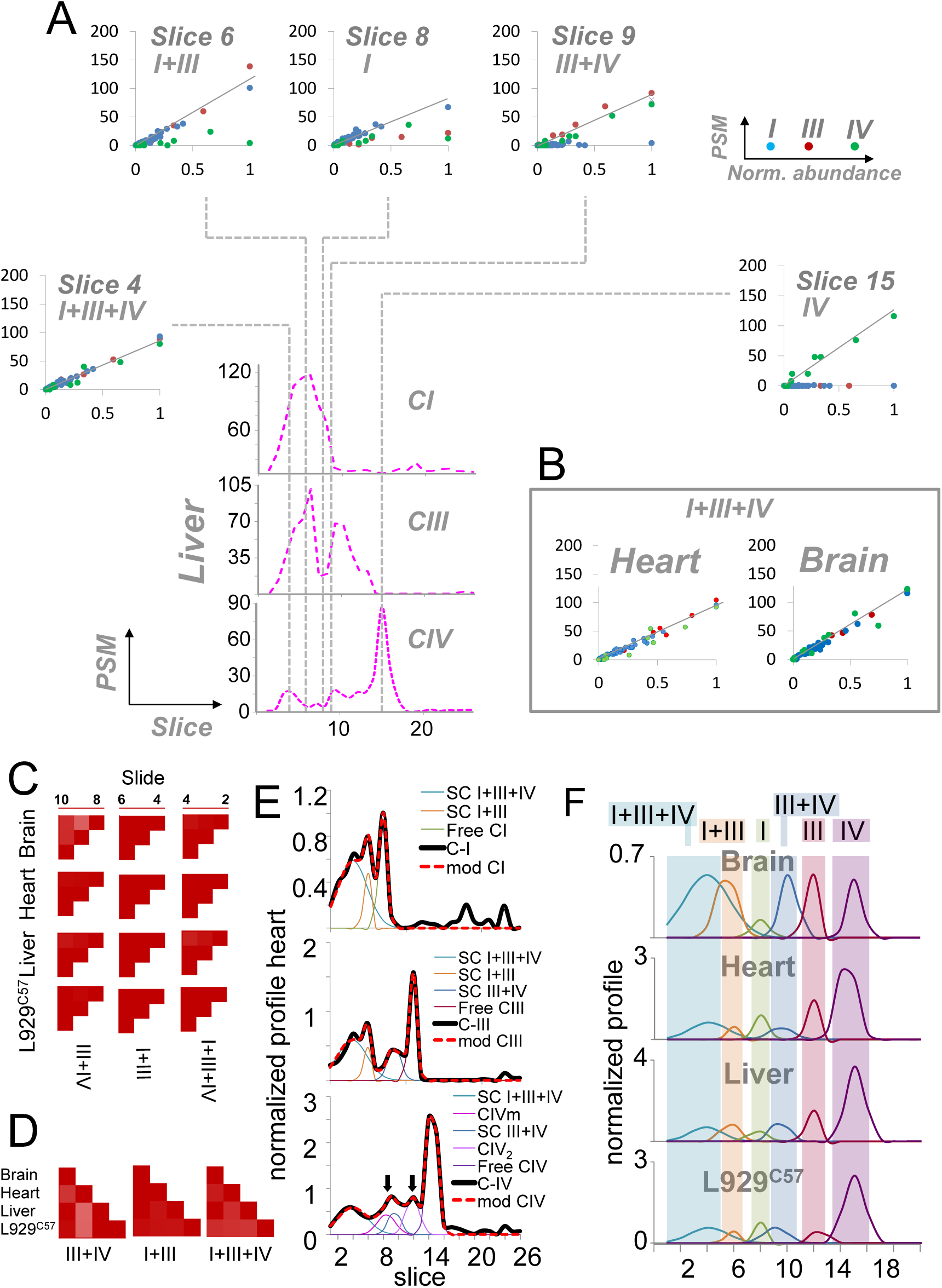
Blue-DiS evidence of the formation of OXPHOS supercomplexes. A, For each complex, the number of protein PSMs was plotted against normalized protein abundances (see Methods), showing that the relative proportions of proteins from CI (blue points), CIII (red points) and/or CIV (green points) are constant in specific BNGE slices from liver mitochondria indicating the presence of multimeric structures. Slices 2-4 correspond to a ternary structure (I+III+IV), slices 5-6 and 9-10 to binary structures (I+III and III+IV, respectively) and slices 8 and 15 to monomeric forms (I and IV, respectively). **B,** Similar results are obtained in heart and brain mitochondria; for simplicity only one slice with the tertiary structure is shown. **C,** Cross-correlation analysis of abundances of proteins forming part of each one of the supercomplexes, showing that their relative proportions are tightly maintained across the consecutive slices. **D,** Cross-correlation analysis of abundances of proteins of the indicated supercomplexes across the four mitochondrial types, showing that their relative proportions are also constant independently of the mitochondrial origin. **E,** Normalized profiles of CI, CIII and CIV from heart mitochondria (black line) (see Methods) can be accurately explained as a superimposition (dashed thick red line) of six gaussian peaks corresponding to monomeric and multimeric structures, as indicated. The gaussian deconvolution was performed in a hypothesis-free manner by a simultaneous least-squares fitting of the three normalized profiles adjusting the position, width and height of each peak (thin lines) without any numeric constraint. In the case of CIV two additional peaks (CIV_2_ and CIVm) had to be added to the model in order to explain the normalized profile (arrows). **F,** Gaussian components used to model the normalized profiles of the four mitochondrial types. For simplicity, the two multimeric CIV structures are not represented.

To further validate the predicted structures of OXPHOS molecular assemblies, we performed a Blue-DiS analysis of mitochondria from cultured mutant cell lines that are unable to assemble one or more of the complexes (Diaz et al., 2006; Moreno-Loshuertos et al., 2006; Perales-Clemente et al., 2010). In the cell line ΔCI, which lacked the ND4 subunit from CI and was unable to assemble CI (Moreno-Loshuertos et al., 2006), neither free CI nor SCs I+III_2_ and I+III_2_+IV could be detected (Fig. S3A, B); however, free III_2_ and IV and the SC III_2_+IV were detected at the expected sizes (green rectangles). In the case of the mutant cell line for Cox10 protein (ΔCIV), which lacks assembled CIV, all structures containing CIV were absent (red rectangles in Fig. S3A, B). In addition, the structures containing CI could neither be detected, in agreement with the fact that CIV is needed for the stabilization of CI (Diaz et al., 2006); hence in this mutant only the free III_2_ form was detectable (green rectangles in Fig. S3A, B). Finally, in the Rho 0 cell line, which lacks mitochondrial DNA encoding for different subunits of the respiratory complexes, none of these structures were observed (red rectangles in Fig. S3A). However, CII, which is totally encoded in the nucleus, and a CV form of smaller size (V*) remained detectable in these cells (Fig. S3A).

In summary, the complete size distributions of OXPHOS components could be explained by structures of constant size and composition that were present in each mitochondrial type at different relative proportions (Fig. 2F).

### Respiratory complexes are assembled into supercomplexes at fixed stoichiometries

The notion that SC contained complexes at fixed stoichiometries was reinforced by the finding that the protein composition of SC was conserved in all mitochondrial types (Fig. 2D). We took advantage of the good linear response of quantitative Blue-DiS protein values (Fig. 2A, B; Fig. S2A) to estimate the relative molar stoichiometries of complexes within each slice (Fig. S3C, D, see M&M for detailed explanation). From slices 2 to 6 the relative molar proportion CIII:CI was 2:1, in the four mitochondrial types, implying that the proportion I+III_2_ is constant in the respirasome and in the SC formed by CI and CIII (Fig. S3E). Besides, the relative molar proportions CI:CIV and CIII:CIV were 1:1 and 2:1, respectively, between slices 2 to 4, implying that the respirasome has a composition I+III_2_+IV (Fig. S3E).

To our knowledge, these are the first stoichiometry estimations of OXPHOS complexes made by mass spectrometry, which agree with the structures analysed by cryo-electron microscopy (Gu et al., 2016; Letts et al., 2016; Sousa et al., 2016), and contradict the experimentally unsupported, but generalized tendency to assume that there are multiple types of respirasomes that differed in their increasing content of 1-4 CIV monomers. The reason for the split of the respirasome in several discrete bands in BNGE, corresponding to a broad peak in Blue-Dis analysis, remains to be clarified. In slices 5 and 6, CIV is still detected although at a lower proportion, suggesting that in this area migrate CIV superstructures of unknown composition (Fig. 3E). Slices 7 and 8 are enriched in CI, where it migrates as a free complex, but again with significant presence of CIV in different high molecular weight structures. Finally, the relative proportions of CIII and CIV in bands 9 and 10 are coherent with a stoichiometry 2:1 (SC: III_2_+IV) (Fig. 3E). In summary, unbiased Blue-DiS analysis reinforces the results obtained by other techniques and provide complementary evidence that OXPHOS complexes are arranged into SC at fixed stoichiometries.

**Figure 3.**
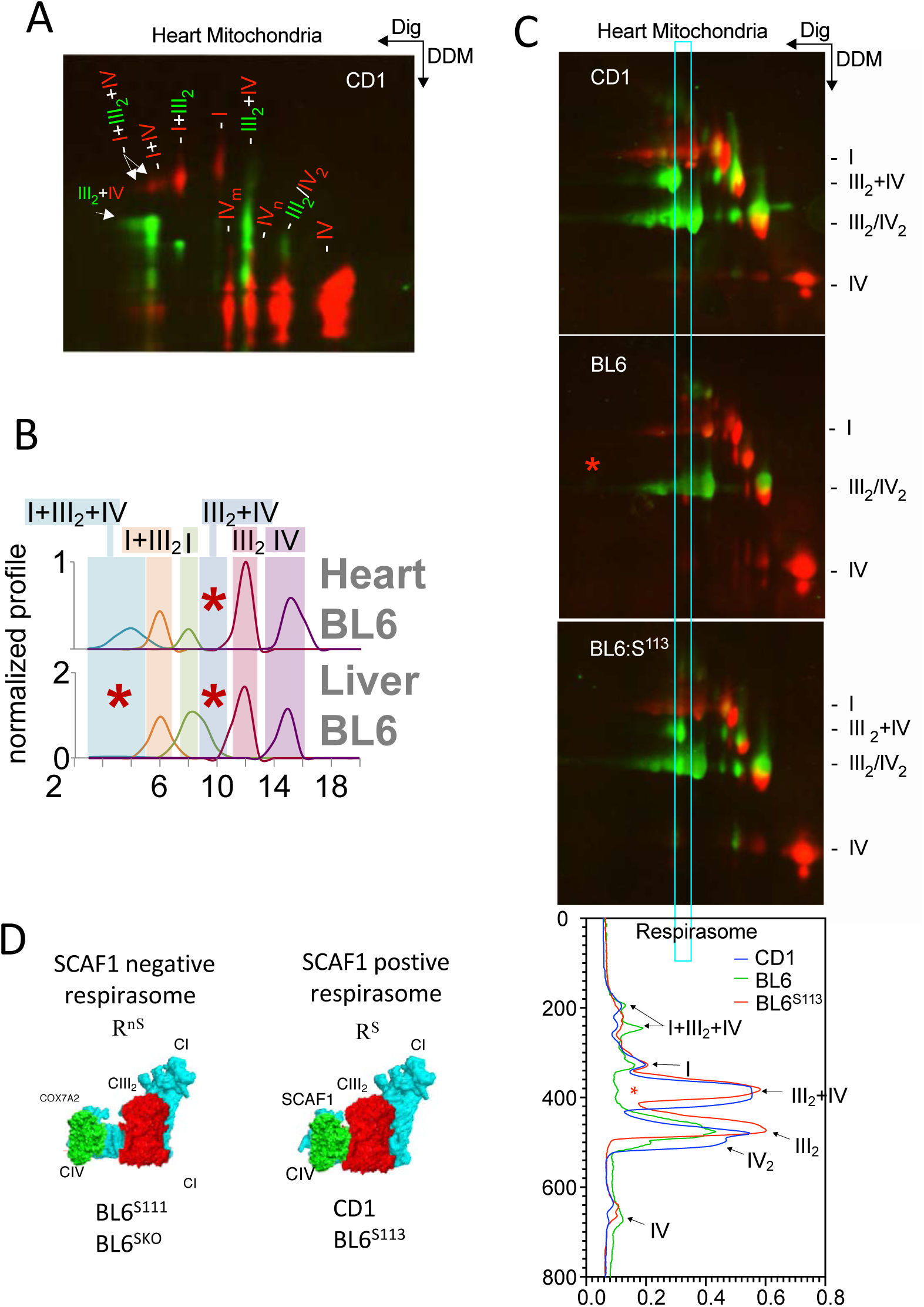
Structural consequences of SCAF1-deficiency in the formation of SCs. **A,** 2DBNGE (Dig/DDM) analysis of heart mitochondria resolving complexes and supercomplexes in the first dimension and disrupting SCs into their component complexes in the second dimension. NDUFA9 immunodetection in red indicate the migration of CI, COI immunodetection in red indicate migration of CIV and CORE2 immunodetection in green indicate migration of CIII. **B,** Gaussian deconvolution of the normalized profiles of heart and liver mitochondria from BL6 mice, where the function of SCAF1 protein is impaired. Red asterisks indicate the position of SC III_2_+IV, which is completely absent in both tissues, and of the respirasome, which is absent in the liver but remains detectable in the heart. **C,** 2DBNGE (Dig/DDM) resolving complexes and supercomplexes in the first dimension and disrupting SCs into their component complexes in the second dimension. Samples from BL6, CD1 and BL6:S^113^ heart are compared highlighting the area of migration of respirasomes, with the different traces at the bottom. Red asterisk indicates absence of the respirasome-derived III_2_+IV only in the BL6 sample indicating that III_2_+IV are not physically linked in SCAF1-deficient respirasomes. NDUFA9 immunodetection in red indicates the migration of CI, COI immunodetection in red indicate migration of CIV and CORE2 immunodetection in green indicate migration of CIII. **D,** representation of the two structurally different respirasomes.

### CIV forms novel supercomplexes

Our complexome analysis predicts that several structures that contain CIV, not yet molecularly characterized, should be present. These are CIV structures above and below SC: III_2_+IV but also some CIV structures of higher molecular weight migration around I+III_2_ and I+III_2_+IV. The existence IV_2_ is well documented, and high molecular weight entities containing only CIV have been described elsewhere (Cogliati et al., 2016). To confirm whether the CIV structures predicted by Blue-DiS analysis are true entities, we performed 2D-BNGE of CD1 heart mitochondria using in the first dimension digitonin as detergent to preserve the integrity of supercomplexes and n-Dodecyl-β-D-maltoside (DDM) in the second dimension to disaggregate supercomplexes into their component complexes. This procedure partially preserves the interaction between CIII and CIV and the IV_2_ (Perales-Clemente et al., 2010). As shown in Fig. 3A this analysis confirmed the presence of IV_2_ and two forms of CIV that migrate immediately faster and slower than SC III_2_+IV and that do not super-assemble with any other respiratory complex (IVm and IVn in Fig. 3A). These three structures account for a significant proportion of the CIV in heart samples. In addition, we detected the presence of low proportions of a putative SC: I+IV_2_, which migrates with SC I+III_2_ and segregates into CI and dimer CIV (Fig. S3F), and of a putative SC I+IV, which migrates between free CI and SC: I+III_2_ (Fig. S3F).

More surprisingly, we found CI and CIV monomer but without CIII that migrate between SC: I+III_2_+IV and SC I+III_2_, very close to the faster migrating respirasome band I+III_2_+IV (Fig. S3F, labelled as I+IV?). The reason why this putative SC migrates with the apparent molecular weight of a respirasome is unclear and may indicate that this structure interacts with a yet to discover mitochondrial inner membrane component. Therefore, 2D-BNGE analysis of heart mitochondria reveals specific supercomplexes that contain CIV, confirming Blue-DiS predictions, and also CIV and CI together and that need to be further investigated. Note that, due to their relatively low abundances, these additional IV structures are fully compatible with the gaussian models of CI and CIV determined from Blue-DiS profiles.

### SCAF1 confers structural attachment between III_2_ and IV within the respirasome

To identify potential interaction partners of specific SC structures from the Blue-DiS analysis, we generated their reference migration profiles *in silico* by combining the SC profiles obtained by deconvolution analysis and performing a correlation search for all identified proteins displaying a similar profile. This analysis led to a natural, hypothesis-free detection of Cox7a2l/SCAF1 as the only candidate showing a significant correlation in all the mitochondrial types with the structures containing CIII and CIV together (ED Fig. 4a left & b). Consistently, deletion of CI inhibited the migration of SCAF1 in the position of the respirasome, but not in the position of SC III_2_+IV, and no traces of SCAF1 were detected in the mutant lacking CIV or in Rho 0 cells (Fig. S4A right). These results obtained by unbiased complexome analysis provide further experimental evidence that this protein regulates the interaction between these complexes, as we postulate in previous reports (Cogliati et al., 2016; Lapuente-Brun et al., 2013).

**Figure 4.**
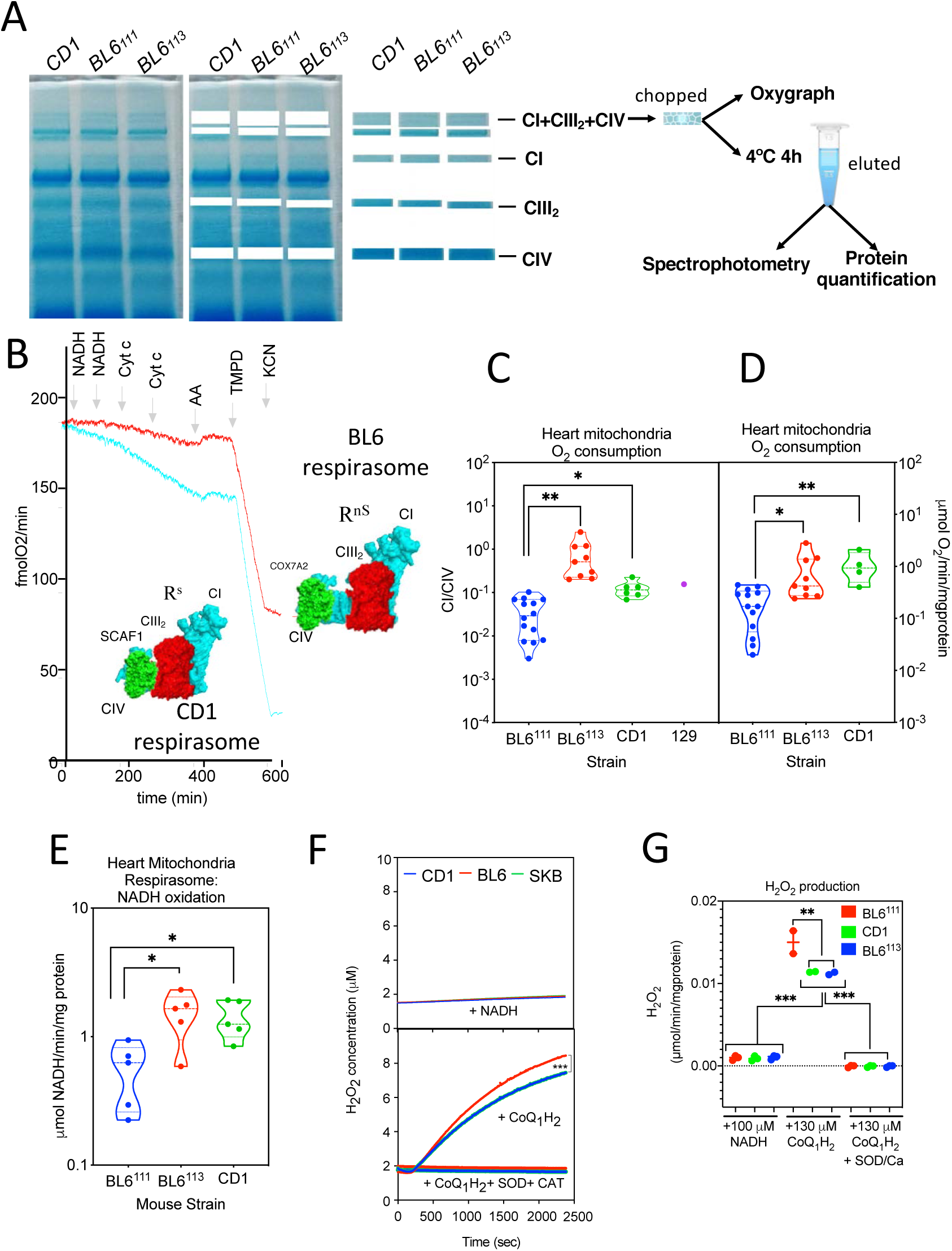
Functional consequences of SCAF1-deficiency in the activity of the respirasome. A, Scheme representing the experimental set up to analyze the function of the respirasomes. **B,** Representative oxygen consumption traces obtained with heart respirasomes excised from the BNGE and derived from either C57BL/6 or CD1 animals. The addition of different components is indicated. **C & D,** NADH-dependent respiration rate normalized by TMPD-respiration rate (**C**), or by mg. of protein (**D**) of heart respirasomes excised from BNGE from the indicated mouse strain and measured in a Clark oxygen electrode. **e,** NADH oxidation rate by heart respirasomes eluted from BNGE excised bands of the indicated mouse strain and measured by spectrophotometry. **F & G,** Representative traces **(F**) and quantitative data **(G)** of H_2_O_2_ production upon NADH oxidation (upper-panel) or CoQH_2_ oxidation (lower panel) by heart respirasomes eluted from BNGE excised bands of the indicated mouse strain and estimated by Amplex Red. When CoQH_2_ oxidation was assayed rotenone was included to prevent the interaction of CoQH_2_ with CI, and the assay was performed.

Next, we analyzed by gaussian deconvolution the Blue-DiS profiles of mitochondria prepared from SCAFI^111^ heart and liver C57BL/6JOlaHsd mitochondria. As shown in Fig. 3B, in BL6 liver the peaks corresponding to the respirasome and the SCs III_2_+IV disappeared, while the rest of structures, including SCs I+III_2_ and the III_2_ peaks, were maintained. However, while SC III_2_+IV was neither observed in BL6 heart, the presence of the respirasome remained clearly detectable in this tissue (Fig. 3B). Therefore, BL6 and CD1 respirasomes are present in heart mitochondria, but differ in the absence of SCAF1 in the former.

We reasoned that if the role of SCAF1 is to physically link CIII and CIV, and this function is maintained in the respirasome, then mitochondria harboring SCAF1^113^ and SCAF1^111^ should have different types of respirasomes differing structurally in the interaction between CIII and CIV. To experimentally evaluate this hypothesis, we took advantage of the fact that upon 2D-BNGE analysis the second dimension used to disaggregate supercomplexes (which employs DDM) does retain significant amounts of the III_2_+IV association, allowing us to discern whether CIII and CIV are physically linked or not in the respirasome in the absence of functional SCAF1. We performed 2D-BNGE with liver mitochondria derived from CD1, C57BL/6JOlaHsd mice and also from C57BL/6JOlaHsd harboring the wild type version of SCAF1 (SCAF1^113^) generated in our laboratory (Cogliati et al., 2016). This approach confirmed again that heart mitochondria from BL6 mice does not assemble SC III_2_+IV (Fig. 3C) and clearly demonstrated that the III_2_+IV structure remains within the respirasomes containing functional SCAF1 (CD1 and SCAF1^113^) but not in those with the non-functional form (BL6^111^) (Fig. 3C). In conclusion, this analysis demonstrates that SCAF1^+^ and SCAF1^-^ respirasomes are structurally different (Fig. 3D).

### SCAF1 enhances respiratory performance and reduce ROS production by the respirasome

We then inquired if the structural difference linked to the presence of an active SCAF1 form in the respirasome has any impact on their respiratory capacity. We first evaluated the suitability to perform these kinetic analyses by confirming in isolated individual complexes (I, III_2_ and IV) that it is possible to measure the rate of oxidation and reduction of their respective donor/acceptor of electrons and that they generate the expected stoichiometries. All the rates of the individual complexes were similar regardless the strain of origin, with the exception of a mild reduction in CIII activity in BL6^113^ (Fig. S4C-E). Next, we excised BNGE bands containing SC I+III_2_+IV (respirasome) from SCAF1^113^ (CD1, BL6^113^ and 129 mice) and SCAF1^111^ (C57BL/6J mice) heart mitochondria and determined their respiratory capacity using NADH as substrate in a Clark electrode (Acín-Pérez et al., 2008) and their NADH oxidation capacity by spectrophotometry (Fig. 4A). For normalization purposes we determined the respiration capacity induced by TMPD in the same samples after inhibition with antimycin and rotenone; we also normalized by the protein content determined after eluting the respirasome from the gel. We found that both SCAF1^+^ and SCAF1^-^-containing respirasomes were able to perform NADH-dependent respiration without the need of adding either external CoQ or Cyt c (Fig. 4B). Interestingly, the respiratory capacity of the respirasome was much lower than that of CIV-monomer and NADH oxidation was also decreased in comparison to that of CI. These results suggest, in agreement with a recent observation on amphipol-stabilized SC:I+III_2_, that superassembly of respiratory complexes modulate their activity (Letts et al., 2019). Moreover, the rate of respiration by SCAF1^+^ respirasomes of different sources was about one order of magnitude higher than that of the SCAF1^-^ ones regardless of whether they are normalized by CIV-dependent respiration (Fig. 4C) or by protein content (Fig. 4D). In agreement with this observation, the rate of NADH oxidization by SCAF1^+^ respirasomes was between 3 and 4 times higher (Fig. 4E).

Lenaz’s laboratory showed by reconstruction in liposomes that the association between CI and CIII into SCs reduces the production of ROS (Maranzana et al., 2013). This observation was validated in vivo by others (Lopez-Fabuel et al., 2016),(Huertas et al., 2017). The availability of respirasomes with different degree of interaction between III_2_ and IV allowed us to investigate whether this interaction impacts on the production of ROS. For this purpose, we incubated the BNGE-excised respirasome bands from CD1, BL6^S111^ and BL6^S113^ heart mitochondria preparations in the presence of NADH and Amplex-red to monitor ROS production. We found no differences in the level of ROS produced by the respirasomes from any of the investigated strains (Fig. 4F, G). Considering that the rate of respiration and NADH oxidation of SCAF1^111^ respirasome is lower than its wild type counterpart, we calculated that the respirasome harbouring the mutant version of SCAF1 derived 0,185% of NADH electrons to ROS, whereas CD1 and SCAF1^113^ derived 0,064 % and 0,067%, respectively. Next, we repeated the experiment but substituting NADH by CoQ_1_H_2_ to donate electrons to CIII and in the presence of rotenone to block CI interaction with CoQ. Under these conditions, which mimic a stress situation with an abnormal rise in CoQH_2_, SCAF1^-^ respirasomes produced significantly more ROS (Fig. 4F, G). In all cases, ROS production was fully quenchable by the addition of superoxide dismutase and catalase (Fig. 4F, G). Therefore, SCAF1^+^ respirasomes produced less ROS than SCAF1^-^ ones.

In summary, although SCAF1 can be dispensable for the formation of respirasomes it confers structural attachment between complexes III_2_ and IV within the respirasomes providing more stability, significantly better functional performance and lower ROS production.

### Respiratory supercomplexes are unstable upon mitochondrial membrane disruption

The existence of two structurally distinct respirasomes was described recently in ovine(Letts et al., 2016) and confirmed later in bovine models (Sousa et al., 2016). One form of respirasome has complexes III_2_ and IV tightly attached (named *tight respirasome*), while the other is characterized by an increased distance between them (named *loose respirasome*). Our observation that SCAF1 determines the interaction between III_2_ and IV within the respirasome is reminiscent of the existence of tight and loose respirasomes. Noticeably, both ovine and bovine SCAF1 sequences match the mouse wild-type and functional 113 version.

Very intriguingly, Letts and co-workers observed that the proportion between both respirasome forms in a given preparation is not stable. In fact, after incubation at 4° C the tight respirasomes were transformed into the loose ones (Letts et al., 2016). We wondered whether this phenomenon could be observed by BNGE analysis. Thus, we compared CD1-liver mitochondria (which harbors the functional SCAF1 form), maintained several hours in the refrigerator, before and after solubilization with digitonin. To our surprise, both SCs I+III_2_+IV and III_2_+IV, but not SC I+III_2_, disappeared in the samples maintained at 4 °C only if they were previously digitonized (Fig. 5A). In addition, SCAF1 migrated as a bulk with the free CIV in the digitonized preparations (Fig. 5A, B). Furthermore, 2D-BNGE/SDS-PAGE analysis revealed that the form of SCAF1 that migrated with free CIV (marked with an asterisk) had a smaller molecular weight than the canonical form (Fig. 5C). Proteomic analysis demonstrated that in slices corresponding to SC I+III_2_+IV, IV_2_ and IV, a peptide spanning the sequence SSVTAYDYSGK (originated from the corresponding SCAF1-derived tryptic peptide LTSSVTAYDYSGK) was unequivocally identified exclusively in the non-fresh samples (Fig. 5E and Fig. S5A). This sequence was mapped into the N-terminus region of the protein, which contains the CIII-interacting domain(Cogliati et al., 2016) (underlined sequence in Fig. 5D). Further quantitative analysis revealed the coexistence of the intact and proteolyzed forms of the peptide bound to CIV in the respirasome, while SC: III_2_+IV contained only the intact form (Fig. 5E). These results indicate that SCAF1 suffers a partial proteolytic processing that disrupts SC: III_2_+IV, destabilizes the respirasome and parallels the loss of CIV-containing SCs. In silico analysis of the processing site in SCAF1 revealed a putative cleavage site for calpain-1 (Fig. 5D). In agreement with this prediction the use of calpain inhibitors was sufficient to prevent the cleavage of SCAF1 (Fig. 5F) and also to preserve the integrity of CIV-containing SC after digitonin treatment (Fig. 5G). When similar experiments were repeated with mitochondrial samples purified from CD1-heart we also observed the relocation of SCAF1 with free CIV, but the reduction in the amount of CIV-containing SC was, although evident, much milder (Fig. 5H). All these results indicate that SC are unstable or can shift from tight (e.g. kinetically active) to loose (e.g. kinetically restrained) forms when mitochondrial membrane integrity is compromised, likely as a consequence of the release of proteases, such as calpain-1, and lipases that gain access to the inner mitochondrial membrane and to the respiratory complexes.

**Figure 5.**
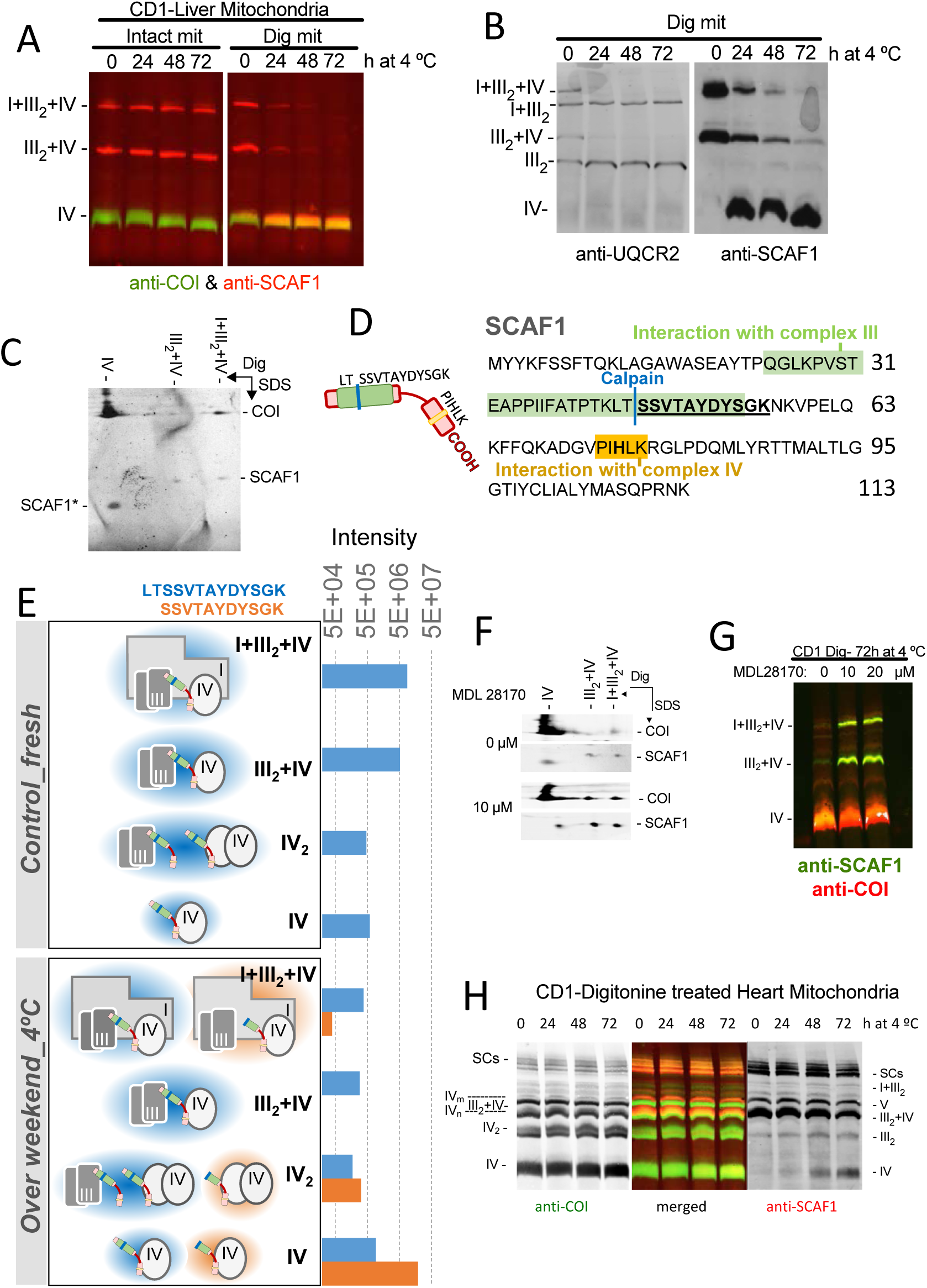
Supercomplexes are unstable upon mitochondrial membrane disruption. A, BNGE resolving complexes and SCs from CD1 liver from intact or digitonized mitochondria preincubated during the indicated time at 4 °C, and probed with the indicated antibody. **B,** BNGE resolving complexes and SCs from CD1 liver from digitonized mitochondria preincubated during the indicated time at 4 °C, and probed with the indicated antibody. **C,** 2D-BNGE/PAGE resolving complexes and SCs in the 1st dimension and protein components in the 2nd dimension from CD1 liver digitonized preparation preincubated 72 h at 4 °C showing that SCAF1 is processed (SCAF1*). The membrane was immunoblotted with the indicated antibodies. **D,** Structure of SCAF1 sequence, mapping CIII-(in green), and CIV-(in yellow) interacting regions. The predicted calpain-1 processing site is indicated with a blue line. **E,** Quantitative analysis of the SCAF1-derived tryptic (LTSSVTAYDYSGK, in blue), and calpain 1-processed (SSVTAYDYSGK, in orange) peptides. Both peptides were quantified in the BNGE slices corresponding to SCs I+III_2_+IV and III_2_+IV and to IV_2_, and to CIV in fresh or in 4°C-incubated liver mitochondria-enriched fractions. The calpain 1-processed peptide was only detected in the non-fresh preparations, attached to CIV and IV_2_ and also to the respirasome. A structural interpretation of these results is presented at the left; the blue and orange shadows indicate whether the SCAF1 peptide is tryptic or processed, respectively. **F,** 2D BNGE/PAGE showing that the proteolytic cleavage of SCAF1 can be prevented by inhibition of Calpain-1. **G,** BNGE/PAGE analysis of liver mitochondria showing that the stability of the respirasome and the SC III2+IV is preserved after digitonization in the presence of a calpain-1 inhibitor. **H,** BNGE profile for CIV (COI-red) and SCAF1 (green) in heart samples maintained at 4°C after digitonization during the indicated time.

It remains to be elucidated whether SCAF1 processing by calpain-1 is just an in vitro artifact or a biologically relevant mechanism regulating SCs turnover dynamics. However, the observation that respiratory complexes become unstable when the mitochondrial membrane integrity is disrupted, raises serious concerns on the interpretation of the various published experimental approaches used to investigate the function of the SCs. They include sub-mitochondrial particles, where functional measurements were performed at 32 °C (Blaza et al., 2014; Fedor and Hirst, 2018); isolated mitochondria, which were incubated at 37 °C (Guarás et al., 2016; Lapuente-Brun et al., 2013); and partially isolated respirasomes (this report). To further study this issue, we subjected digitonized and intact mitochondria from liver (Fig. S5B) and heart (Fig. S5C) preparations to prolonged incubations at different temperatures. Liver digitonized mitochondria completely lost the respirasome and the SC III_2_+IV, after 72 hours at 4 °C, and all SCs, including I+III_2_ after 3h at 37 °C (Fig. 5A and Fig. S5B). Respirasomes from heart digitonized mitochondria were more stable than those from liver at 4 °C (72h) but were completely lost after 3h at 37 °C; at this temperature the SC III_2_+IV and even the CIV monomer disappeared (Fig. S5C). In contrast, all of these structures remained stable in intact organelles (Fig. S5B, C). More worrying is the fact that the stability of both heart and liver SCs is not maintained if digitonized mitochondria are incubated at 37 °C for just one hour (Fig. S5D). All these observations call to caution when performing experiments in disrupted mitochondria. Very importantly, the differential stability of SC in intact and broken mitochondria may well be the cause of the apparently discrepant results regarding the physiological role of respiratory supercomplexes between different groups.

Our unexpected results prompted us to evaluate if BNGE purified respirasomes were stable enough during functional studies to trust the conclusions presented above. To address this issue, we kept the first-dimension gel for 1 or 3 hours at 37°C after running the BNGE to inquire if the respirasomes are preserved. After the incubation period we ran a second-dimension electrophoresis also in the presence of digitonin to determine the intactness of structures, since complexes and SCs are expected to migrate forming a diagonal unless some associations are disrupted during incubation. Thus, after 1- or 3-hour incubation at 37 °C the respirasome and any other detected complex or SCs remained substantially intact apart from comet tail-like shape of the spots due to diffusion in the 1^st^ dimension gel during the long period of incubation (Fig. S5E). Moreover, 2D-BNGE analysis, using DDM in the second dimension to dissociate the respirasomes, confirmed the stability of the interaction between III_2_ and IV within the respirasome (Fig. S5F). Therefore, BNGE allows the isolation of the respirasome maintaining its structure to perform functional studies, validating our functional observations.

### Superassembly of OXPHOS complexes influences the delivery of electrons and the respiratory capacity within the electron transport chain

The lack of stability of the respiratory supercomplexes upon disruption of mitochondrial membranes raises serious concerns on the interpretation of recent experiments where recombinant AOX was added to a preparation of sub-mitochondrial particles (Fedor and Hirst, 2018). The data obtained suggested that the interaction of CI and CIII in supercomplexes does not modify the delivery of electrons to the alternative oxidase AOX by CoQ. These results led to the conclusion that there is a unique CoQ pool equally accessible to CI and CII within mitochondrial inner membrane. Given the observation that the native structure of supercomplexes is preserved in intact mitochondria, we performed similar experiments but expressing AOX in the mitochondria of different cell models: a) wild-type cells expressing AOX (CIV^WT^AOX and E9AOX); b) cells lacking CIV which retain CI and SCI+III_2_ superassembly due to the presence of AOX (CIV^KO^AOX); c) cells that lack CIII which preserve CI due to AOX expression but CI cannot form supercomplexes with CIII (CIII^KO^AOX). AOX expressed in mammalian mitochondria is known to be functional and can recycle oxidized CoQ in cells lacking of mtDNA (Perales-Clemente et al., 2008) or in cells lacking CIII or CIV (Guarás et al., 2016). We described elsewhere that the expression of AOX prevent the degradation of CI in the absence of CIII or CIV (Guarás et al., 2016) (Fig. S6A). We found that the level of AOX activity was similar between the different cell lines (Fig. S6B). Following similar reasoning to that of Fan & Hirst, the delivery of electrons from CI or CII to AOX should be independent of superassembly in the absence of CIII. However, if the superassembly between CI and CIII has any effect on the activity of CI, in the absence of CIV a change in the delivery of electrons from CI to AOX should be detected (Fig. 6A). In the two mutant cells we detected measurable CI-AOX and CII-AOX respirations, while CIV-dependent respiration could not be recorded (Fig. 6B). Strikingly, CII-AOX dependent respiration was insensitive to the presence of CIII, while CI-AOX dependent respiration was significantly lower when CIII is present (Fig. 6B). This is true despite a higher NADH oxidation capacity of CI in CIV^KO^AOX cells (Fig. 6C). We also reasoned that if a CoQ pool was shared between CII and CI, addition of succinate would outcompete with glutamate + malate-based AOX respiration (Fig. 6D). We found that in either the wild type or in the CIV^KO^ cells, all expressing AOX, the addition of succinate significantly increased the oxygen consumption over that achieved by CI substrates (Fig. S6C, D and Fig. 6E). On the contrary, in CIII^KO^ AOX cells the addition of succinate was unable to increase the AOX dependent oxygen consumption over that reached by CI substrates (Fig.S6D and Fig. 6E). These results suggest that when CI is not superassembled CoQ exist in a unique pool, whereas its superassembly triggers the formation of two partially differentiated pools.

**Figure 6.**
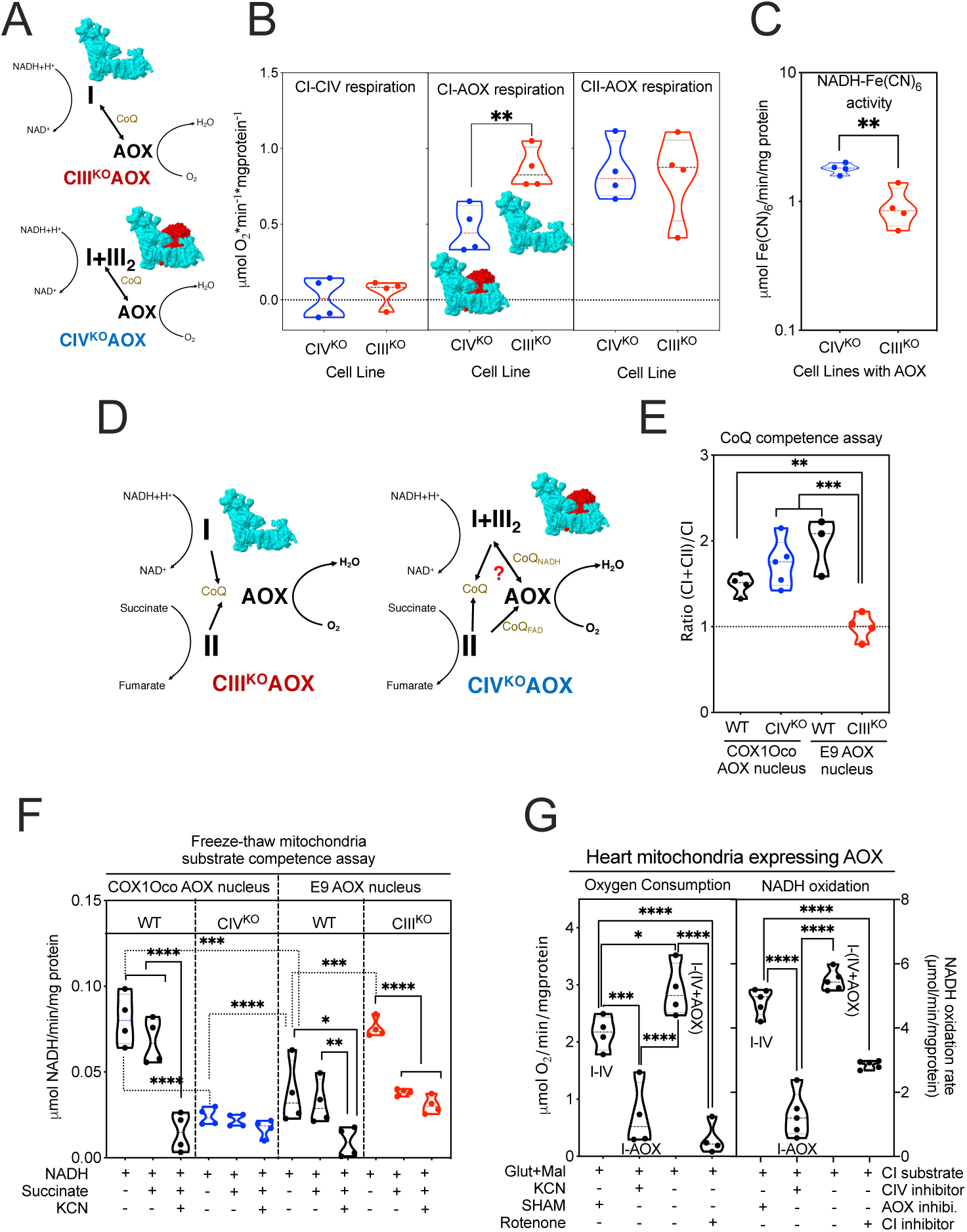
The superassembly between complexes I and III modulates the activity of CI and functionally segments the CoQ pool. A, Scheme representing the differential flux of electrons to AOX from the indicated mutant cell line. **B,** Impact of the presence of CIII in the delivery of electrons from CI and CII to AOX. **C,** DPI sensitive NADH oxidation capacity of the mitochondrial preparation from the indicated cell line. **D,** Scheme representing the simultaneous flux of electrons form either CI or CII to AOX from the indicated mutant cell line. **E,** Estimation of the impact of the simultaneous addition of substrates for CII (succinate) on the CI-dependent respiration with CI substrates (glutamate and malate) in the presence or absence of CIII. **F,** Impact of CIII and CIV superassembly on the maximum respiration capacity with CII (succinate) or CI (glutamate and malate) substrates vs. both substrates addsed simultaneously. **G**, Proportion of the maximum respiration archivable by CI substrates in the presence or absence of CI and CIII superassembly. **H,** Analysis of the flux of electrons from NADH and CI or succinate and CII to AOX in the indicated freeze-thaw mitochondrial from wild type cells or mutant cells lacking CIV or CIII all expressing AOX. **I,** Flux of electrons from NADH and CI to AOX or CIV in intact heart mitochondria expressing AOX and monitored by oxygen consumption (left panel) or autofluorescence of NADH (right panel).

We then measured directly NADH or succinate dependent respiration (Fig. 6F, G) and NADH oxidation (Fig. 6H) in mitochondrial membranes permeabilized by freeze & thawing in a period of 4 min to prevent disruption of supercomplexes, and isolating mitochondria from wild type, CIV^KO^ and CIII^KO^ cell models, all expressing AOX. This analysis offered a number of interesting observations: (i) In the presence of CIII none of the substrates (NADH or succinate) was able to reach the oxygen consumption levels obtained when both substrates were added simultaneously (Fig 6F, G). This indicates that the delivery pathway of electrons to CoQ and AOX is partially different for each substrate. (ii) NADH oxidation of the two wild-type cells lines is different, being the E9 nuclear background significantly lower (≈50%) (Fig. 6H). We described elsewhere that this is due to the presence of a missense mutation in COI that reduce the activity of CIV, and hence, respiration (Acin-Perez, 2003), a fact that can also be observed in ED Fig.5C and D. (iii) The addition of succinate did not impact on the oxidation of NADH in any wild type cell line while the inhibition of CIV by KCN almost completely abolished it in wild type cells (Fig. 6H). (iv) The ablation of CIV dramatically reduced the rate of oxidation of NADH to levels equivalent to those caused by KCN (Fig. 6H). (iv) The ablation of CIII significantly increased the NADH oxidation with respect to the levels of their isogenic wild type cells -which harbor a mutation in COI-(Fig. 6H). This result parallels what we have observed measuring oxygen consumption in intact mitochondria (Fig. S6D), and fully confirms that the dependence of CIV activity limits the NADH oxidation capacity in E9 derived cells. (v) In the absence of CIII but not of CIV, the addition of succinate significantly reduced the NADH oxidation, which in these cells is fully dependent of AOX (Fig. 6H). All these results demonstrate that the superassembly of CI with CIII, either in SCs or in the respirasome, impacts dramatically in the NADH oxidation capacity of CI by modifying the delivery of electrons to CIV or AOX. Moreover, they indicate that CI and CII can potentially outcompete for delivering electrons to AOX, a phenomenon that is minimized by the presence of CIII. Our results with AOX expression substantially disagree with those published by Fedor and Hirst (Fedor and Hirst, 2018). The reason for this discrepancy is likely due to the unnoticed disruption of the supercomplexes under their experimental set up, as explained above.

There are, however, additional experimental differences between our analyses that may contribute to those discrepancies. Fedor et al. utilized mitochondria extracted from hearts as starting material for their experiments (Fedor and Hirst, 2018). In order to minimize the impact of the mitochondrial source, we generated a novel mouse model expressing AOX in the heart and muscle mitochondria (see material and methods for details) and repeated the experiments with purified wild type muscle and heart mitochondria expressing AOX. We first performed the analysis of respiration in intact mitochondria fed with CI substrates (Glut+Mal) and assessing the distribution of electrons either to CIV or AOX by using specific inhibitors for each enzyme (Fig. 6I, left panel). As expected, more than 70% of CI-dependent respiration was sensitive to CIV inhibition, with a significant respiration mediated by AOX (Fig. 6I, left panel). We confirmed that this assay was specific for CI by monitoring NADH oxidation in intact mitochondria by measuring changes in NADH auto-fluorescence during the assay in the absence or in the presence of rotenone (Fig. 6I, right panel). We additionally performed experiments in freeze-thawed mitochondria prepared from hearts expressing AOX and confirmed again that NADH-linked respiration was completely inhibited by KCN, whereas CII-AOX respiration was maintained (Fig. S6E). Monitoring NADH absorbance allowed us to confirm that KCN completely abolished NADH oxidation, in contrast to SHAM, which did not have any significant effect (Fig. S6F). Similar results were obtained with mitochondrial preparations from muscle expressing AOX (Fig. S6G, H). In conclusion, our results based on AOX expression could be reproduced using different sources of mitochondria and demonstrate that CI supercomplex assembly affects the delivery of electrons to AOX.

## Discussion

In this report we provide strong experimental support demonstrating that the superassembly of respiratory complexes into supercomplexes induce novel characteristics that deeply impact on the structure, kinetics and efficiency of the respiratory chain. We demonstrate it using isolated supercomplexes, mitochondrial preparations and cell cultures, and also using animal models. We also provide experimental data that explains why previously published results apparently contradict the main conclusion of this paper.

The main findings supporting our conclusion can be summarized as follows: 1) The absence of functional SCAF1 in mice causes a reduction in exercise performance both in males and females and compromise growth under food restriction in males. 2) The absence of functional SCAF1 eliminates the physical interaction between CIII and CIV in the SC III_2_+IV or within the respirasome (I+III_2_+IV). 3) In the absence of functional SCAF1, the respirasome can be assembled in some tissues despite the fact that the direct interaction between CIII and CIV is absent. 4) The absence of SCAF1 destabilizes the respirasome, reduces both oxygen consumption and NADH oxidation and increases ROS production. 5) The interaction between CI and CIII decreases the rates of NADH oxidation and respiration by AOX, implying the existence of a partially segmented CoQ pool driven by the superassembly of CI and CIII.

Overall, the more relevant conclusion of our work is that the formation of respiratory supercomplexes modify the kinetics and the flux of electrons occurring with non-superassembled complexes. In one side, the role of SCAF1 in the respirasome becomes now clear. After strong discrepancies, there is a general consensus that SCAF1 is required for the superassembly of SC:III_2_+IV and to provide stability to the respirasome (Cogliati et al., 2016; Lobo-Jarne et al., 2018). There is also agreement in that SCAF1 function is lost in the mutated form of SCAF1 that naturally arise in the C57BL/6 sub-strains (Enríquez, 2019; Lapuente-Brun et al., 2013). In spite of that, BL6 heart mitochondria still assemble bona-fide respirasomes, which are structurally different, less stable and functionally impaired. These findings clarify the role of SCAF1 in the respirasome and the relevance of the interaction between CIII and CIV. On the other side, the expression of AOX in cells and in mice and the ability to genetically control the free or superassembled status of CI, allowed us to confirm that the delivery of electrons from CI but not CII to AOX or CIV is asymmetric only if the formation of supercomplex I+III_2_ is occurring. These results confirm that superassembly of CI and CIII allow a partially segregation of the CoQ pool. Interestingly, a very recent paper from the group of Sazanov studying the kinetic properties of NADH to Cyt c electron transfer of the isolated supercomplex I+III_2_ also found that CoQ trapping within isolated respiratory supercomplex I+III_2_ limits CI turnover (Letts et al., 2019).

In addition to the major findings, we exhaustive re-evaluated the mitochondrial BNGE profiles from a variety of murine sources and analyzed the stability of the sample under several conditions using advanced data-independent proteomics. This analysis provided relevant new knowledge to better interpret BNGE analysis. In one hand, the Blue-DiS technology allowed us to confirm, from a true hypothesis-free perspective, the presence, composition and stoichiometry of the OXPHOS CI, CIII, CIV and CV in free and superassembled forms. Thus, while the protein composition of these complexes is maintained, the proportion of OXPHOS supercomplexes is tissue-specific. Moreover we demonstrate that regardless of the different positions where the respirasome migrate, it maintains a constant stoichiometry of I+III_2_+IV, a result that is in full agreement with the different cryo-electron-microscopy derived structures published to date (Gu et al., 2016; Letts et al., 2016; Sousa et al., 2016; Wu et al., 2016). This estimation stresses the necessity to correct the experimentally unsupported tendency to assume that the different bands of the respirasome contain increasing amounts of complex IV and to define the band with the faster migration rate as the I+III_2_+IV respirasome and those with a higher apparent molecular weight as successive megacomplexes (Lobo-Jarne et al., 2018). It is advisable to restrict the use of the term megacomplex to define the association between SCs (Guo et al., 2017). Nevertheless, it is remarkable that the respirasome migrate persistently in multiple bands, a phenomenon that require further investigation of the potential partners. The Blue-DiS analysis also confirmed, from a completely unbiased perspective, that SCAF1 is the main assembly factor which is present in SC containing CIII and CIV. On the other hand, we describe unequivocally the existence of novel SCs: I+IV that co-migrate between the free CI and SC: I+III_2_ and I+IV_2_ that co-migrate with SC: I+III_2_; and the co-migration of CI with CIV in the area of the respirasomes. CI and CIV associations could be predicted from the evidence obtained by cryo-electron-microscopy that the interaction between CI and CIII_2_ is independent from that between CI and CIV (Letts et al., 2016; Sousa et al., 2016). SC I+VI_2_ was however unexpected. This novel SC, as well as some of the already known associations may represent true SCs or partially disassembled elements from larger complexes. In any case, their characterization in the BNGE is of major relevance because they can lead to inaccurate interpretations. This may be the case of the proposed novel fast migrating respirasome (I+III_2_+IV) that comigrate with I+III_2_, despite the different molecular weight, solely based on the fact the CI, CIII and CIV co-migrate in the same band (Sun et al., 2016). The co-migration of I+III_2_ with I+IV_2_ is a more plausible explanation, since they have a very similar MW. The co-migration CI with CIV in the area of the respirasomes, well above SC: I+III_2_, does not contain IV dimers and the added molecular weight of CI and CIV does not justify its apparent MW. The fact that 2D-BNGE/DDM splits CI and CIV suggest as the more plausible explanation that this co-migration imply the presence of a SC I+IV that interact with an unknown partner.

This additional SC may again flaw our interpretation of the BNGE gels. Thus, when the gels are performed with a steep gradient or short run (as it is the case of the more used pre-casted native gels) the SCs I+III_2_+IV (the faster migrating band) and this particular SC: (I+IV)* may overlap (as a potential example see (Lobo-Jarne et al., 2018)) leading to the wrong interpretation solely based on the position of CIV that this band correspond to a respirasome.

A third important discovery reported here is that the respirasome and the SC: III_2_+IV, and at less extent SC I+III_2_, are unstable after disruption of the mitochondrial membranes even if preserved at 4 °C. This previously unnoticed phenomenon raises concerns in the interpretation of experiments aimed to measure the proportion and function of SCs when mitochondrial membrane integrity is not preserved. More strikingly, the loss of respirasome and SC: III_2_+IV is paralleled by the specific proteolytic processing of SCAF1 by calpain-1 that causes the processed form of SCAF1 to migrate together with CIV. If this cleavage is physiologically relevant or not need to be further investigated, but at this point it explains discrepant results in the literature that considered that SCAF1 as a shorter protein that acts as a mere isoform of CIV (Zhang et al., 2016). By the same token it may explain the unexpected presence of variable amounts of SCAF1 associated with free CIV. Our observation is also coherent with the description of two respirasome structures by several groups (Letts et al., 2016; Sousa et al., 2016) that differ in the connection between III_2_ and IV that is either present (tight respirasome) or absent (loose respirasome). Our finding also explains the reported conversion of the tight respirasome into the loose one with time (Letts et al., 2016). Therefore, these observations largely clarify the investigation of the functional role of the respiratory supercomplexes using BNGE.

In summary, in this report we show that superassembly between respiratory complexes substantially enhances the respiratory capacity of the electron transport chain while minimizing ROS production. Superassembly establishes a segmented CoQ pool through the association between CI and CIII, while SCAF1 plays a critical role in the III_2_+IV interaction affecting the structure and kinetic properties of the respirasome. Hence, SCAF1 is a key factor in the regulation of the energy metabolism, optimizing efficiency under high energy demands or restricted availability of nutrients.

## Acknowledgments

This study was supported by MINECO: SAF2015-65633-R, MCIU: RTI2018-099357-B-I00, CIBERFES (CB16/10/00282) and HFSP (RGP0016/2018) to JAE; MINECO-BIO2015-67580-P, and PGC2018-097019-B-I00, ISCIII-IPT13/0001, ISCIII-SGEFI/FEDER, ProteoRed) the Fundació MaratóTV3 (grant 122/C/2015) and “la Caixa” Banking Foundation (project code HR17-00247) to JV. The CNIC is supported by the Ministry of Economy, Industry and Competitiveness (MEIC) and the Pro-CNIC Foundation, and is a Severo Ochoa Center of Excellence (MINECO award SEV-2015-0505).

## Author contributions

EC, JV & JAE: conceived and designed the analysis, EC, ML-L, FG-M-M. and JS-C performed the proteomic analysis. SC, AG, PH-A RA-P, YM and MC-A: performed the BNGE and the respirasomes functional analysis. SC, MC-A, J.R.H and R.A.C performed the *in vivo* experiments. EC, SC, ML-L, JV, PH-A, JV and JAE interpreted the results. EC, JV and JAE wrote the paper.

## STAR METHODS

### LEAD CONTACT AND MATERIALS AVAILABILITY

Request of information and material should be made to José Antonio Enriquez. Mouse and cell lines generated in this study are available upon request for a non-commercial use under a Material Transfer agreement.

### EXPERIMENTAL MODEL AND SUBJECT DETAILS

#### Experimental Models

This study used mouse and cellular models which were generated in out laboratory.

#### Mouse generation

***1)*** C57BL/6JOlaHsd mice with the functional version of SCAF1 were generated as previously described(Cogliati et al., 2016). C57BL/6JOlaHsd mice knock out for SCAF1 were generated by microinjection of ES cells knock-out in the first alleles from EuMMCR repository in a C57BL/6JOlaHsd blastocysts. Further, the blastocysts were implanted in a pseudo pregnant female C57BL/6JOlaHsd. 2) AOX expressing mice were generated by *genOway* by targeted insertion of the AOX cDNA within the Rosa26 locus via homologous recombination in embryonic stem cells originally derived from a 129 strain of mouse and injected into C57BL/6J blastocysts and then re-implanted into OF1 pseudo-pregnant, and allowed to develop to term. After removal the neo cassette flanked by FLP sites, the selected knock-in animasl were then sistematically backcrossed for more than 20 generations to C57BL/6JOlaHsd background. The expression of the transgene is dependent upon the Cre recombinase mediated excision of a *Lox*P flanked transcriptional “STOP” cassette upstream the AOX cDNA. For this study we induce the expression of AOX in muscle and heart by breeding the AOX animals with expressing CRE under the ACTA promoter.

#### Mouse experimentation

All animal procedures conformed to EU Directive 86/609/EEC and Recommendation 2007/526/EC regarding the protection of animals used for experimental and other scientific purposes, enforced in Spanish law under Real Decreto 1201/2005. Approval of the different experimental protocols requires the estimation of the adequate sample size as well as the definition of the randomization and blinding criteria. The mice were fed a standard chow diet unless other food regime was required (5K67 LabDiet).

#### Alternative fasting protocol

Starting from weaning, mice undergo to 48h of fasting alternate between 24h of controlled feeding (standard chow diet 5K67LabDiet Rod18-A; LASQCdiet) both with free access to water for 40 days. The weight has been recorded at the end of feeding phase.

#### Maximal incremental running test

Mice was forced to run on a treadmill at 20° slope (LE8700 (76-0303), Treadmill Panlab, Harvard Apparatus). After a minute of acclimatization at 10 cm/s, the speed was increased up to 16 cm/s for 5 minutes and further every 2 minutes by 3cm/s until exhaustion, that was considered to reach when the mouse spent 3s in the back of the rack.

#### Cellular Models

All cell lines were grown in DMEM (GibcoBRL) supplemented with 5% FBS (foetal bovine serum, Gibco BRL). mtDNA-less mouse cells were generated by long term growth of L929 mouse cell line (ATCC CCL-1) in the presence of high concentrations of Ethidium Bromide (EthBr) as previously described(Acin-Perez, 2003). Control cells (control cells) were generated by transformation of ρ°929neo cells by cytoplast fusion using NIH3T3 fibroblasts as mitochondrial donors(Bayona-Bafaluy et al., 2008).

## METHOD DETAILS

### Blue-Native Gel Electrophoresis

Supercomplex levels and compositions were analyzed in isolated mitochondria from different tissues and cells by blue native electrophoresis (BNGE)(Wittig et al., 2006). Mitochondrial proteins from heart tissue were solubilized with 10% digitonin (4g/g) (Sigma D5628) and run on a 3%–13% gradient Blue Native gel. The gradient gel was prepared in 1.5 mm glass plates using a gradient former connected to a peristaltic pump. After electrophoresis, the gels were further processed for proteomic analysis, western blotting, 2D SDS-PAGE or 2D-BNGE (DDM) analysis. For 2D SDS-PAGE, the first-dimension lanes were excised from the gel and incubated 1h at room temperature in 1% SDS and 1% β-mercaptoethanol and run in a 16.5% second denaturing gel. For 2D-BNGE (DDM) and 2D-BNGE Digitonin, first-dimension lanes were excised from the gel and run in a 3-13% gradient gel in native condition adding 0.02% of DDM to the cathode buffer.

### Immunodetection of complexes and supercomplexes

After BNGE, 2DBNGE/DIG, 2DBNGE/DDM or 2DBNGE/PAGE, proteins were electroblotted onto PVDF transfer membrane (Immobilon-FL, 0.45 µm, Merck millipore, IPFL00010) for 1 h at 100 V in transfer buffer (48 mM Tris, 39 mM glycine, 20 % EtOH). A Mini Trans-Blot Cell system (BioRad) was used. Sea Blocking buffer (Thermo Scientific 37527) or PBS with 5 % BSA was used for 1 hour at room temperature (RT) to avoid non-specific binding of antibodies. For protein detection, antibodies were incubated with the membrane for 2 hours at RT. Secondary antibodies were incubated for 45 minutes at RT. The membrane was washed with PBS 0.1 % Tween-20 for 5 minutes three times between primary and secondary antibodies and after secondary antibodies, the last wash was only PBS. To study supercomplexes assembly, the PVDF membrane was sequentially probed with antibodies Complex I (anti-NDUFA9, Abcam ab14713), Complex IV (anti-COI. Invitrogen 35-8100), Complex III (anti-core2, Proteintech), SCAF1 (anti-COX7A2L, St. John’s laboratory STJ42268. This antibody was generated by immunization of rabbit with KLH conjugated synthetic peptide between 37-65 amino acids from the central region of human COX7A2L. It recognizes an epitope in the common part of full size and processed SCAF1, allowing to visualize the processed SCAF1 migrated with CIV after Calpain processing as in Fig.5c and Fig.5f. This antibody was discontinued and substituted by the anti-COX7A2L, St. John’s laboratory STJ110597 produced with a full length recombinant human CoX7A2L that, in the same experimental conditions does not recognize the calpain-1 processed SCAF1 that need to be identified only by MS. Anti-MIC10 (MINOS1 Novusbio NBP1-91587) and anti-CHCHD3 (Proteintech 25625-1-AP).

### Proteomics by data-independent scanning (DiS) mass spectrometry (MS)

DiS is a data-independent acquisition method that covers all possible fragmentations of precursors in the 400-1100 m/z range in two LC-MS runs and has already been successfully used to study the mitochondrial proteome(Cogliati et al., 2016; Glytsou et al., 2016; Quintana-Cabrera et al., 2018). DiS uses narrow MS/MS windows of 2 m/z, typical of data-dependent acquisition methods, allowing direct peptide identification by database searching and FDR control by using a conventional target/decoy competition strategy, without requiring peptide fragmentation libraries. The Blue-DiS workflow generated a permanent, multi-dimensional, high-resolution time-fragment mass map for all possible precursors present in each BNGE fraction and each mitochondrial sample, from which quantitative protein maps can be straightforwardly obtained with minimal computation. BNGE gels were excised in 26 slices taking as reference some discrete Coomassie stained bands: slice 6 corresponds to a band that mainly contains SC I + III_2_, slice 10 to CV, slice 12 to free CIII_2_, and slice 15 to free CIV. All slices were cut into cubes (2×2 mm), reduced with 10mM DTT (GE Healthcare), alkylated with 55mM iodoacetamide (Sigma-Aldrich) and subjected to a standard overnight in-gel digestion at 37°C with 3 µg of sequencing grade trypsin (Promega, Madison, WI, USA) in 100 mM ammonium bicarbonate, pH 7.8. After desalting with C18 Omix cartridges (Agilent Technologies), the resulting tryptic peptide mixtures were injected onto a C-18 reversed phase (RP) nano-column (75µm I.D. and 50 cm, Acclaim PepMap, Thermo Fisher, San José, CA, USA) using an EASY-nLC 1000 liquid chromatography system (Thermo Fisher, San José, CA, USA) and analysed in a continuous gradient consisting of 8–31% B for 130 min, 50–90% B for 1 min (B= 0.5% formic acid in acetonitrile). Peptides were eluted from the RP nanocolumn at a flow rate of ∼200nl min^−1^ to an emitter nanospray needle for real-time ionization and peptide fragmentation in either a Q-Exactive or a Q-Exactive HF mass spectrometer (Thermo Fisher). Each sample was analysed in two chromatographic runs covering different mass ranges (from 400 to 750 Da, and from 750 to 1,100 Da, respectively). The DiS cycle consisted of 175 sequential HCD MS/MS fragmentation events with 2-Da windows that covered the whole 350 Da range. HCD fragmentation was performed at 30 normalized collisional energy, a resolution of 17,500 and a maximum injection time of 80 ms with the AGC set to a target of 3 ×10^5^ ions. The whole cycle lasted 30 s or less depending on ion intensity during chromatography. Peptide identification was performed using Sequest running under Proteome Discoverer 1.4 (Thermo Fisher Scientific), allowing two missed cleavages, and using 2 Da and 20 p.p.m. precursor and fragment mass tolerances, respectively. Met oxidation and Cys carbamydomethylation were selected as dynamic and static modifications, respectively. FDR for peptide identification was controlled using a separate inverted database and the refined method(Navarro and Vázquez, 2009). Visualization, validation and quantification of MS/MS spectra from specific peptides was performed using Vseq script, as described(Cogliati et al., 2016).

### Protein and complex profiling

Quantitative protein migration profiles from Blue-DiS analyses were obtained by spectral counting, summing up the number of peptide-spectrum matches (PSMs) of all peptides identified for each protein on each slice of the gel. In the case of protein complexes, the total PSMs of the proteins contained in the complex was used for quantification. The normalized BNGE profile of each complex was constructed by calculating the slope from the plot of protein PSMs in a given slice versus protein PSMs in a slice used as reference (the one with the highest number of PSMs). The normalized abundance of each protein within a complex was calculated as the slope from the plot of PSMs of the protein in the different slices versus the PSMs of the most abundant protein from the complex in the same slices.

### Determination of stoichiometry between complexes in SC

We observed that the number of PSMs for each protein, as calculated by DiS, was approximately proportional to the number of tryptic peptides detectable by MS (NOP) (ED Fig. 3C, left). This finding agrees well with the use of the “protein abundance index” by other authors, which normalizes spectral counts by the number of detectable peptides(Ishihama, 2005). This consideration allowed us to calibrate the individual MS response of each protein, generating an estimation of the effective NOP (ED Fig. 3C, right). By plotting, per each complex, the PSMs of the proteins against their effective NOP, we could estimate from the slopes the molar stoichiometries of complexes within each slice (ED Fig. 3d).

### Activity of complexes or respirasomes from BNGE eluted bands

CD1, BL6:S^111^ and BL6:S^113^ heart mitochondria were extracted, processed and run in BN-PAGE as described above. CI, CIII_2_, CIV-monomer or respirasome bands were quickly excised from gels and minced on ice. The grist was immediately resuspended in Medium MAITE and respirasomes eluted by twirling for 4 hours at 4°C. Elution was collected and mixed with the appropriate volume of MAITE + 2.5 mg/ml BSA at 37 °C to reach 1 mL. Reaction was started by adding 100 µM NADH, 130 µM CoQ_1_, reduced DQ 130 µM, oxidized Cyt c 100 µM. NADH, oxidized CoQ_1_ and Cyt c levels were tracked by recording absorbance at 340 nm, 289 nm and 550 nm, respectively for 240 sec in a UV/VISJASCO spectrophotometer. Optimal absorbance values were calculated by titration of each reactive in MAITE + 2.5 mg/ml BSA. DQ maximal absorbance peak could not be estimated due to the consequent turbidity of its dissolution in an aqueous environment. Baseline NADH oxidation and CoQ_1_ reduction from the same elution was recorded after addition of 1 µM rotenone. Baseline Cyt c reduction from the same elution was recorded after addition of 2.5 µM antimycin A. Baseline Cyt c oxidation from the same elution was recorded after addition of 1 mM KCN.

### Oxygen consumption by BNGE respirasome bands

CD1, BL6:S^111^ and BL6:S^113^ heart mitochondria were extracted, processed and run in BN-PAGE as described above. CIV-monomer or respirasome bands were quickly excised from gels and minced on ice. The grist was immediately resuspended in MAITE + 2.5 mg/ml BSA at 37°C and introduced in an Oxytherm System S1/MINI. Reaction was started by adding 100 µM reduced Cyt c for CIV-monomer or 100 µM NADH for respirasomes. Oxygen levels were tracked for at least 180 seconds and baseline oxygen consumption was recorded after addition of 1 mM KCN for CIV-monomer or 1 µM rotenone for respirasomes.

### H_2_O_2_ production by respirasomes from BNGE eluted bands

CD1, BL6:S^111^ and BL6:S^113^ heart mitochondria were extracted, processed and run in BN-PAGE as described above. Respirasome bands were quickly excised from gels and minced on ice. The grist was immediately resuspended in Medium MAITE and respirasomes eluted by twirling for 4 hours at 4°C. Elution was collected and mixed with Amplex Red solution (Molecular probes), following manufacturer’s instructions. Reaction was started by adding 100 µM NADH or 100 µM CoQH_2_+1µM rotenone. Amplex Red levels were tracked by recording fluorescence at 540/590 nm (exc/emm) for 2400 sec in a Fluoroskan Ascent fluorimeter (Thermo Labsystems). Baseline Amplex Red fluorescence from the same elution was recorded after addition of 5U/mL of SOD and Catalase.

### NADH oxidation monitoring in intact mitochondria by autofluorescence

AOX expressing mice heart mitochondria were extracted as described above and immediately resuspended in medium MAITE at 4°C. Mitochondria were mixed with the appropriate volume of MAITE+ 2.5 mg/ml BSA at 37 °C to reach 100 µl. Reaction was started by adding 5 mM Glu + 5 mM Mal ± 1 mM KCN, 5 mM SHAM or ± 1 µM rotenone. NADH levels were tracked by recording autofluorescence at excitation/emission of 340/475 nm for 20 min in a Fluoroskan Ascent fluorimeter (Thermo Labsystems)

### Oxygen consumption measurement

AOX expressing heart mitochondria of mitochondria from AOX expressing cell lines were extracted as described above and immediately resuspended in medium MAITE at 4°C. To permeabilize mitochondrial membranes, mitochondria were subjected to a freeze-thaw step. Intact or freeze-thawed mitochondria were mixed with the appropriate volume of MAITE + 2.5 mg/ml BSA at 37 °C and placed in an Oxytherm System S1/MINI. Reaction was started by adding 5 mM Glu + 5 mM Mal ± 10 mM succinate ± 1 mM KCN, 5 mM SHAM or ± 1 µM rotenone. Permeabilized mitochondria were provided with 100 µM NADH or 100 µM CoQ_1_H_2_. Baseline levels were recorded after addition of 1µM rotenone or 5 mM SHAM.

### Activity of complexes in freeze-thaw mitochondria

AOX expressing heart mitochondria of mitochondria from AOX expressing cell lines were extracted as described above. Mitochondria were immediately resuspended in medium MAITE and subjected to a freeze-thaw step. Freeze-thaw mitochondria were mixed with the appropriate volume of MAITE + 2.5 mg/ml BSA at 37 °C to reach 1 mL. Reaction was started by adding 100 µM NADH and in some cases 1 mM Fe(CN)_6_, 1 µM rotenone + 1 µM antimycin A, 100 µM succinate or 1 mM KCN was added to the reaction mixture. Since the accessibility of CoQ-analogs could be differential across the different AOX-cell models (i.e. with or without supercomplexes), we performed DPI-sensitive NADH:Fe(CN)_6_ activity (i.e. an activity for FMN in CI) which allowed us to quantify CI-content. NADH and Fe(CN)_6_ levels were tracked by recording absorbance at 340 nm and 412 nm, respectively for 240 sec in a UV/VISJASCO spectrophotometer. Baseline NADH consumption from the same sample was recorded after addition of 1 µM rotenone. Baseline Fe(CN)_6_ absorbance from the same sample was recorded after addition of 5 µM diphenyleneiodium (DPI).

### QUANTIFICATION AND STATISTICAL ANALYSIS

#### Statistical analysis

Unless specified, statistical analyses and graphics were produced with GraphPad Prism 8 software. Data sets were compared by t-test, ANOVA or non-parametric analysis when corresponded and with p-values adjusted for multiple test. Differences were considered statistically significant at P values below 0.05. \**p-value* < 0.05; ** *p-value* < 0.01; *** *p-value* < 0.001; **** *p-value* <0.0001. All results are presented as mean ± SD or mean ± SEM.

## DATA AND CODE AVAILABILITY

Some of the datasets supporting the current study have not been deposited in a public repository yet, but will be done when possible, meanwhile they are available from the corresponding author on request.

