## Supplementary material for "Functional role of respiratory supercomplexes in mice: segmentation of the Q_pool_ and SCAF1": Suplementary Information

**Supplementary figures**:

Figure S1.- Frequency of death caused by the adaptation to restricted food supply in the indicated mouse strains. Related to figures 1 and 2.

Figure S2. The mitochondrial respiratory complexes have the same protein composition but a tissue-specific molecular weight distribution. Related to figure 2

Figure S3. -Validation of structures predicted by Blue-DiS analysis. Related to figure 3

Figure S4.- Identification of SCAF1 as a $III_2$+IV assembly factor. Related to figure 4

Figure S5.- Degradation of SCAF1 by calpain-1. Related to Figure 5

Figure S6.- AOX as a tool to investigate the segmentation of the CoQ pool. Related to figure 6

**Supplementary Tables and Datasets**:

Supplementary Table 1.- Cross-correlation analysis of protein abundances in BNGE slices within each one of the protein complexes.

Supplementary Table 2.- Cross-correlation analysis of protein abundances in representative BNGE slices of each complex across the four mitochondrial types

Dataset 1.- List of Peptide-Spectrum Matches in BNGE slices in all biological models

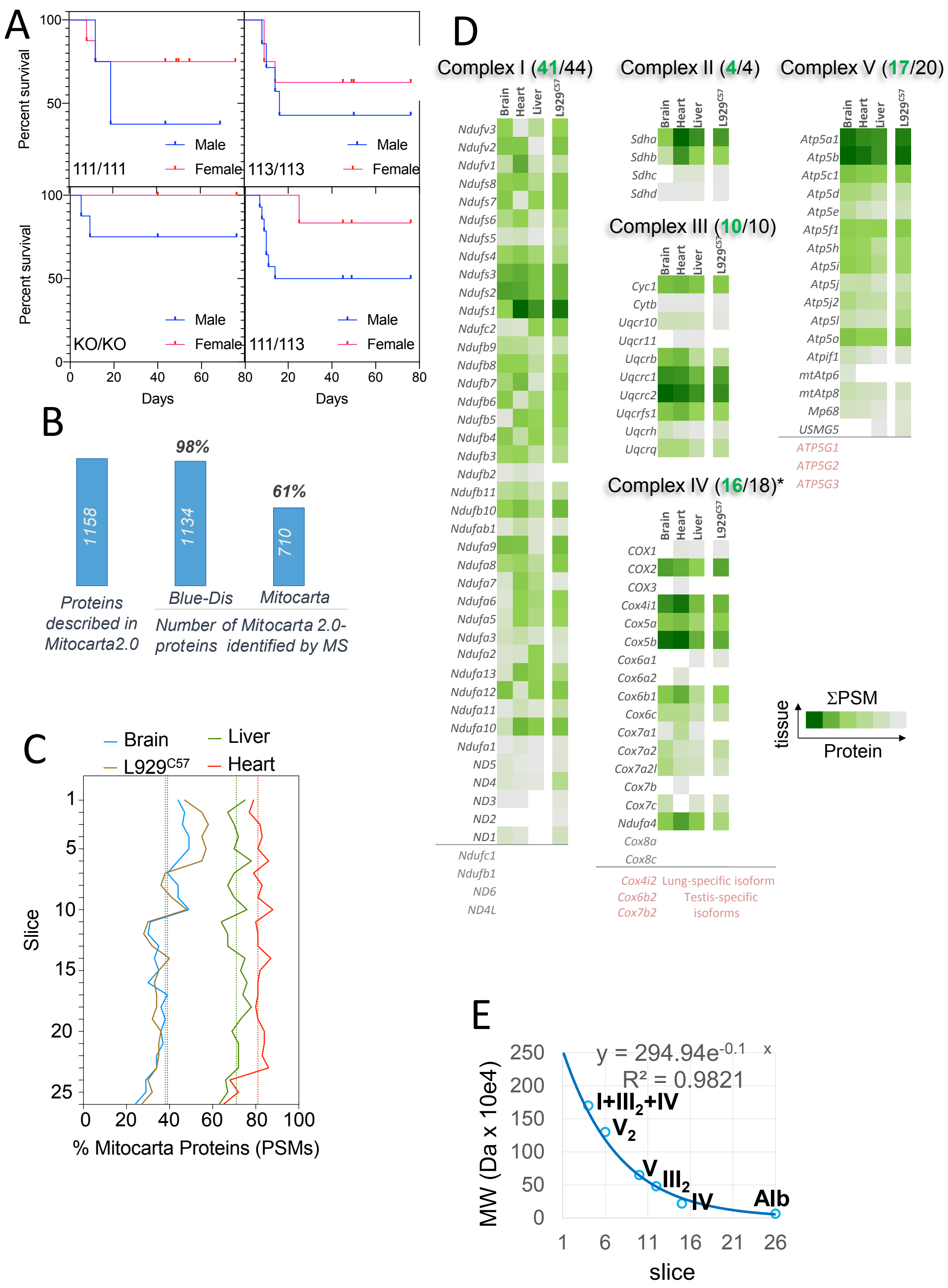


**Figure S1.- Frequency of death caused by the adaptation to restricted food supply in the indicated mouse strains. A,** Protein coverage obtained by Blue-DiS**. B**, Total coverage of mitochondrial proteins reached by Blue-DiS. **C**, Coverage of mitochondrial proteins along BNGE slices in the four mitochondrial sources. **D,** Coverage of structural proteins belonging to CI, CII, CIII, CIV and CV. **E**, Correspondence between apparent molecular weight and slice number in BNGE migration profiles.

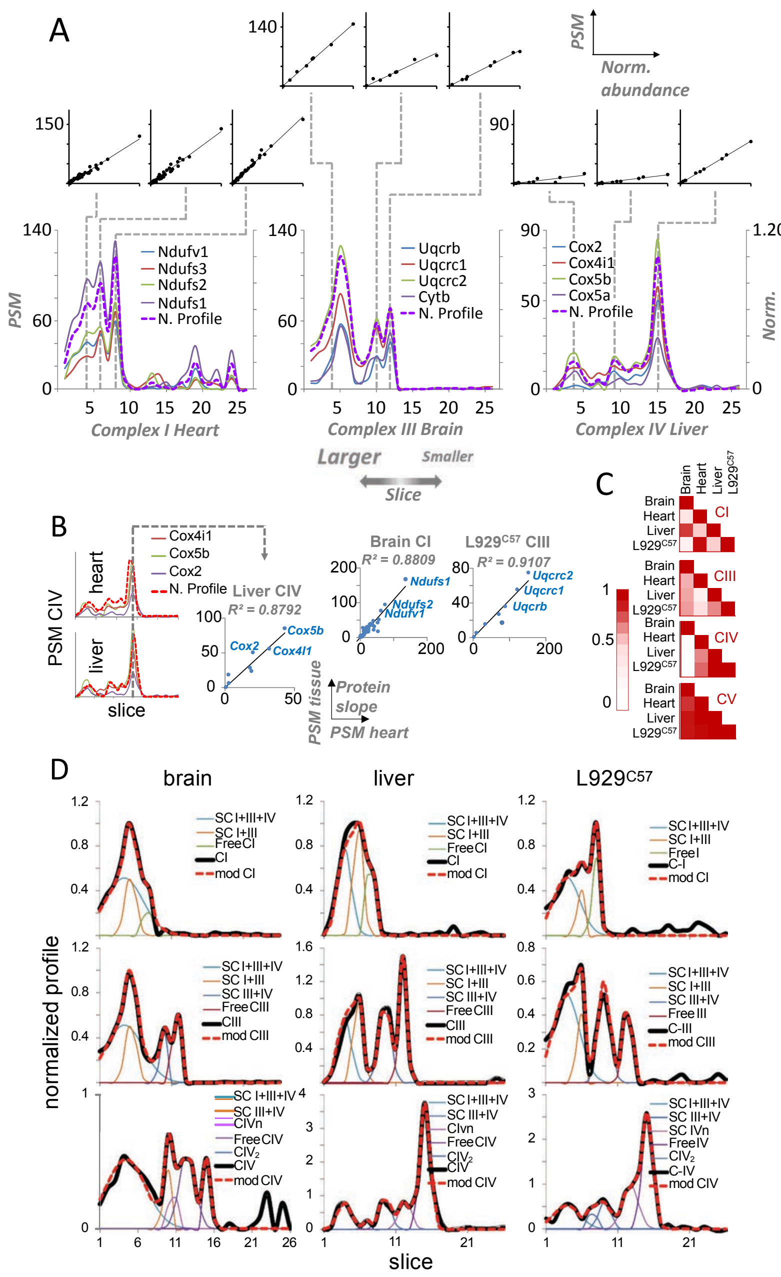


**Figure S2. The mitochondrial respiratory complexes have the same protein composition but a tissue-specific molecular weight distribution**. **A,** Cross-correlation analysis of protein abundances of different BNGE slices within the same complex and mitochondrial source. Correlation coefficients are calculated against the normalized protein profiles of each complex (see Material and Methods). The strong correlations reveal that the complexes have constant protein proportions. **B,** Cross-correlation analysis of protein abundances of the most representative peak from each complex across the four mitochondrial sources. The figure shows representative results revealing a very high correlation between the abundances of CIV proteins in liver (left), CI proteins in brain (middle), and CIII proteins in brain (right), and the same proteins in heart. These results demonstrate that these complexes are composed by proteins in the same proportions in the four mitochondrial types. **C,** Cross-correlation analysis of the normalized profiles of each complex across the four mitochondrial types. The results show that, with the exception of CV, the molecular weight distribution of each complex is not maintained across the mitochondrial types, revealing a tissue-specific size distribution. **D**, Profiles of CI, CIII and CIV-containing complexes from brain, liver and L929 (black thick lines) and their gaussian deconvolution model (red thick dashed lines). The gaussian components are indicated by thin lines, as in Fig. 2D.

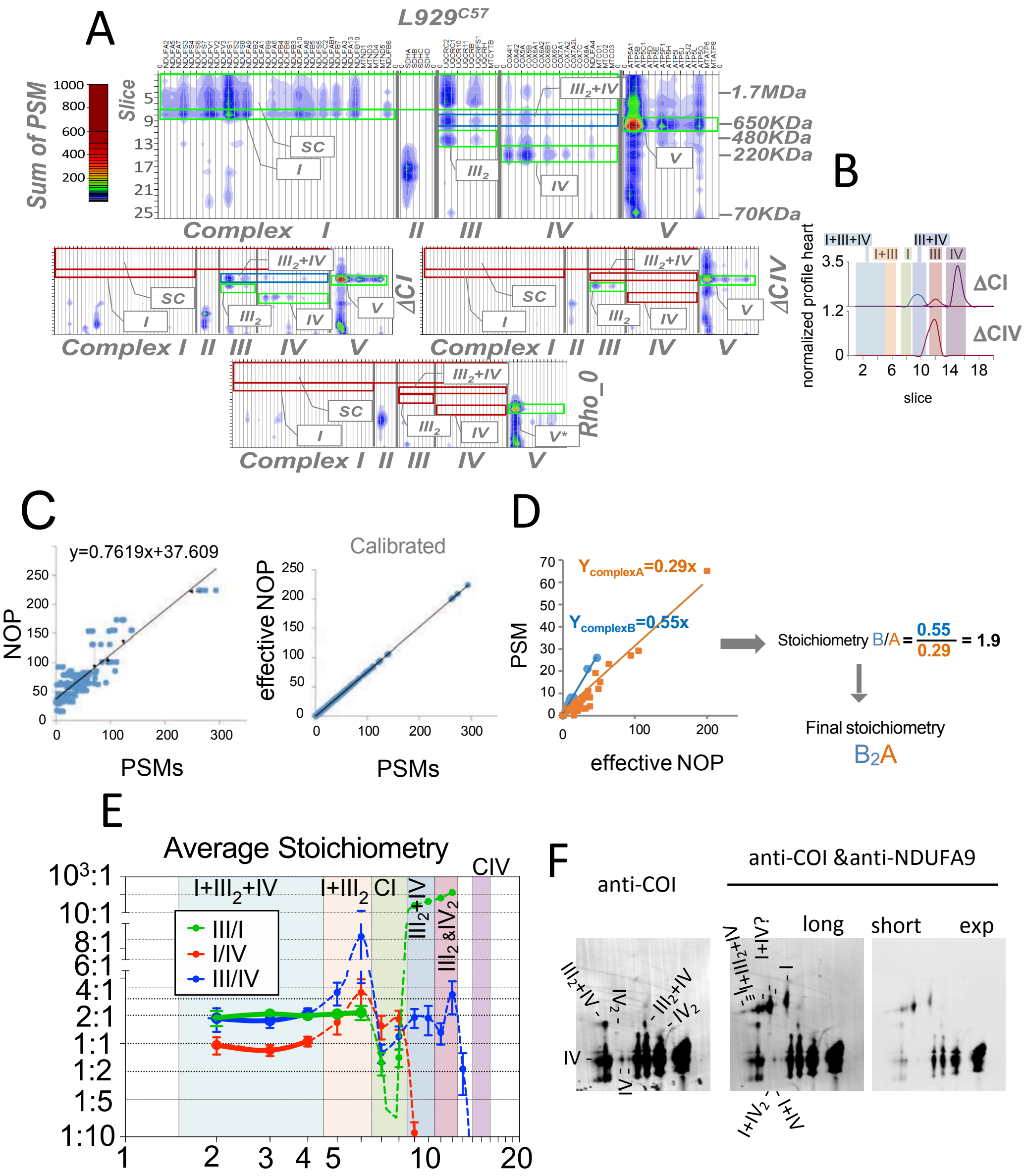


**Figure S3. Validation of structures predicted by Blue-DiS analysis. A,** Contour plots representing the abundance (in number of PSM) for CI, CII, CIII, CIV and CV proteins in control (L929C57), CI- (ΔC1) and CIV- (ΔC4) deficient cells or in cells lacking mtDNA (Rho 0). **B,** Gaussian deconvolution models of the normalized profiles of mitochondria from the two mutant cell lines ΔC1 (which cannot assemble CI) and ΔC4 (which are unable to assemble CIV). **C to E,** Determination of the stoichiometry of the complexes**. C**, Calibration of individual MS protein response by estimating the effective number of tryptic peptides detectable by MS (effective NOP). The number of tryptic peptides (NOP) is plotted against the number of PSM of each protein, producing an almost straight line (left). This line is used to calculate the effective NOP of each protein, defined as the NOP the protein would have to generate the observed PSM (right). **D**, The stoichiometry of OXPHOS complexes is estimated by plotting the observed PSM per protein from each complex against their effective NOP; the stoichiometry between two complexes is estimated as the ratio of the slopes from each one of them. **E,** OXPHOS complexes are arranged into supercomplexes at a fixed stoichiometry. The graph shows the mean ± SEM of the stoichiometry calculated in the four mitochondrial types. The large points joined with thick lines highlight the stoichiometries that can be accurately calculated without interference from other structures. **F,** novel CIV containing supercomplexes, 2DBNGE (Dig/DDM) resolving complexes and supercomplexes in the first dimension and disrupting SCs into their component complexes in the second dimension. COI immunodetection was performed first to indicate migration of CIV containing structures, NDUFA9 immunodetection was performed subsequently indicate migration of CI containing structures. This gel corresponds to that shown in main figure 5E where the immunodetection of CORE2 to localize the migration of CIII is also included.

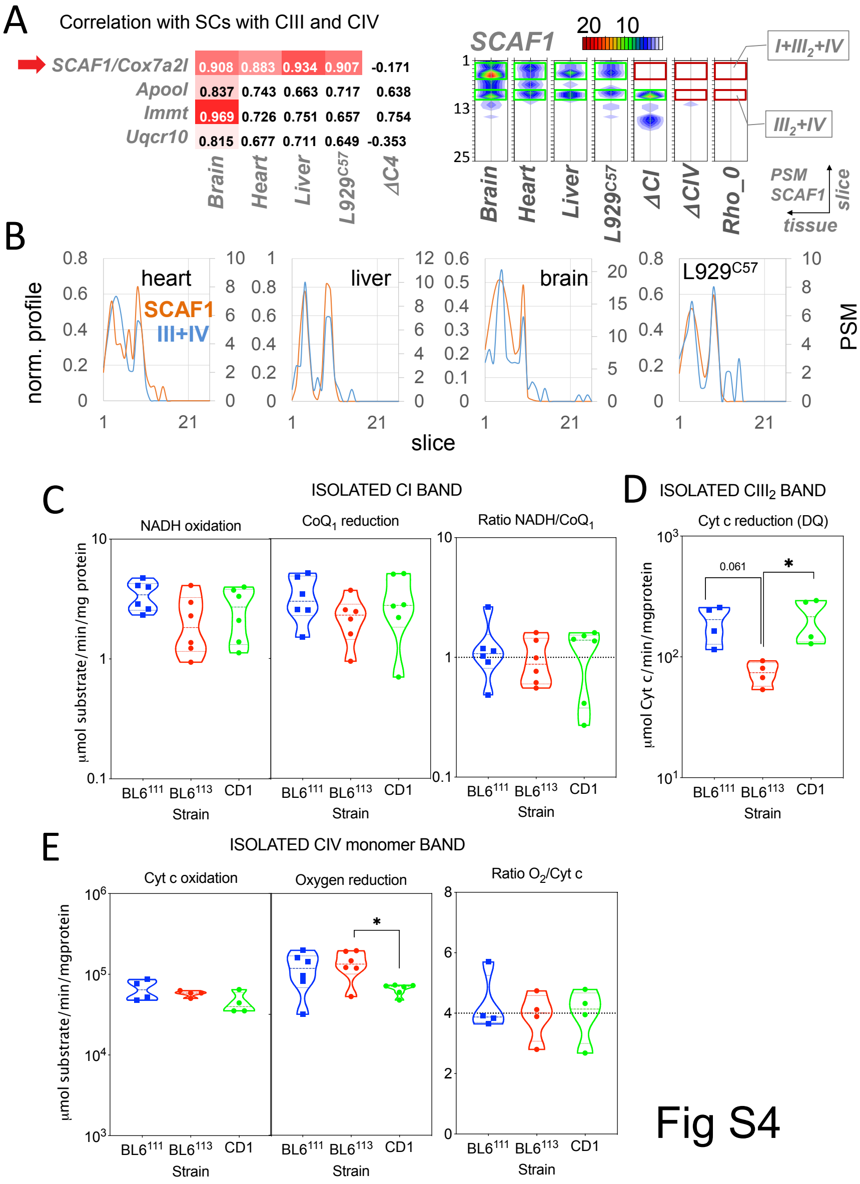


**Figure S4. Identification of SCAF1 as a III$_2$+IV assembly factor. A- left**, SCAF1 is the only protein whose Blue-DiS profile has a significant correlation with the in-silico profiles of SCs containing CIII and CIV together (SC I+III$_2$+IV and III$_2$+IV) in the four mitochondrial types (left). The table shows the four proteins with the highest correlation scores among all proteins detected by Blue-Dis. **A- right,** Contour plots showing the quantitative distribution of SCAF1 in the four mitochondrial types. The migration position for SCs I+III$_2$+IV and III$_2$+IV are indicted in green or red squares. **B**, Agreement between Blue-DiS profiles of SCAF1 and those of SCs containing CIII and CIV together in the indicated mitochondrial types. **C to E**, Characterization of functional activity of the BNGE isolated complexes**. C-left,** NADH-oxidation vs CoQ$_1$ reduction rates by CI eluted from BNGE excised bands from heart of the indicated mouse strain, measured in a spectrophotometer. **C-right**, molar ratio between NADH and CoQ$_1$ electron transfer. **D**, Cytochrome c oxidation rate by isolated CIII$_2$ from the same sources, feed with decylubiquinol (DQ) and measured by spectrophotometry. **E-left,** Cytochrome c oxidation vs oxygen reduction rates of isolated monomer complex IV eluted from BNGE excised bands from the indicated mouse strain, measured by spectrophotometry and in a Clark oxygen electrode, respectively. **E-right**, molar ratio between cytochrome c and oxygen electron transfer.

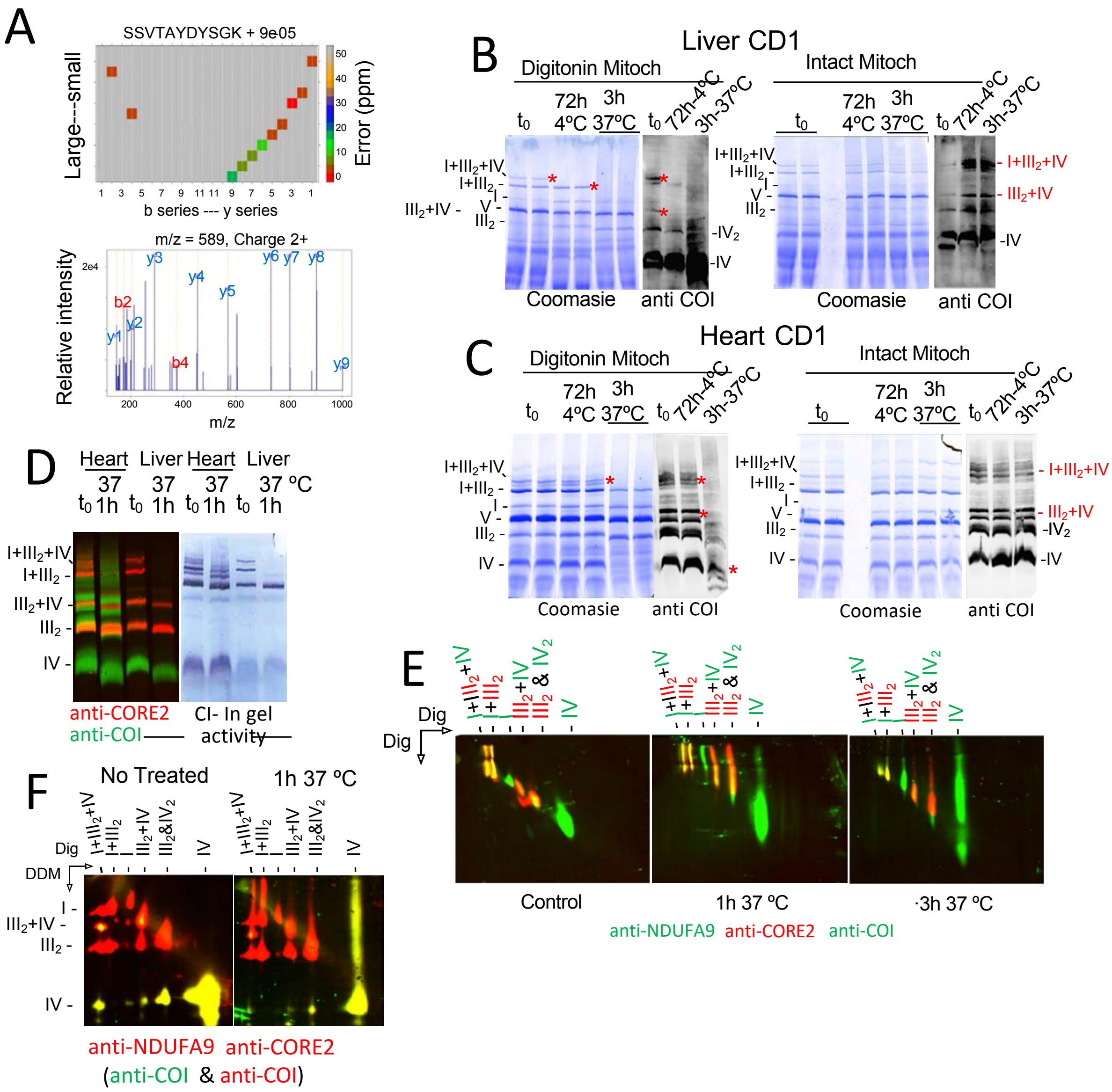


**Figure S5. Degradation of SCAF1 by calpain-1. A,** Vseq analysis validate the identification by MS/MS of the SCAF1-derived processed peptide (non-triptic) spanning the sequence SSVTAYDYSGK. **B-D,** BNGE analysis of the supercomplex integrity in the indicated samples incubated in the indicated conditions after digitonization or maintained intact. **E,** 2D BNGE Dig/Dig analysis of liver mitochondria maintained in the indicated conditions between the first and the second dimension. **F,** 2D BNGE Dig/DMEM analysis of liver mitochondria maintained in the indicated conditions between the first and the second dimension.

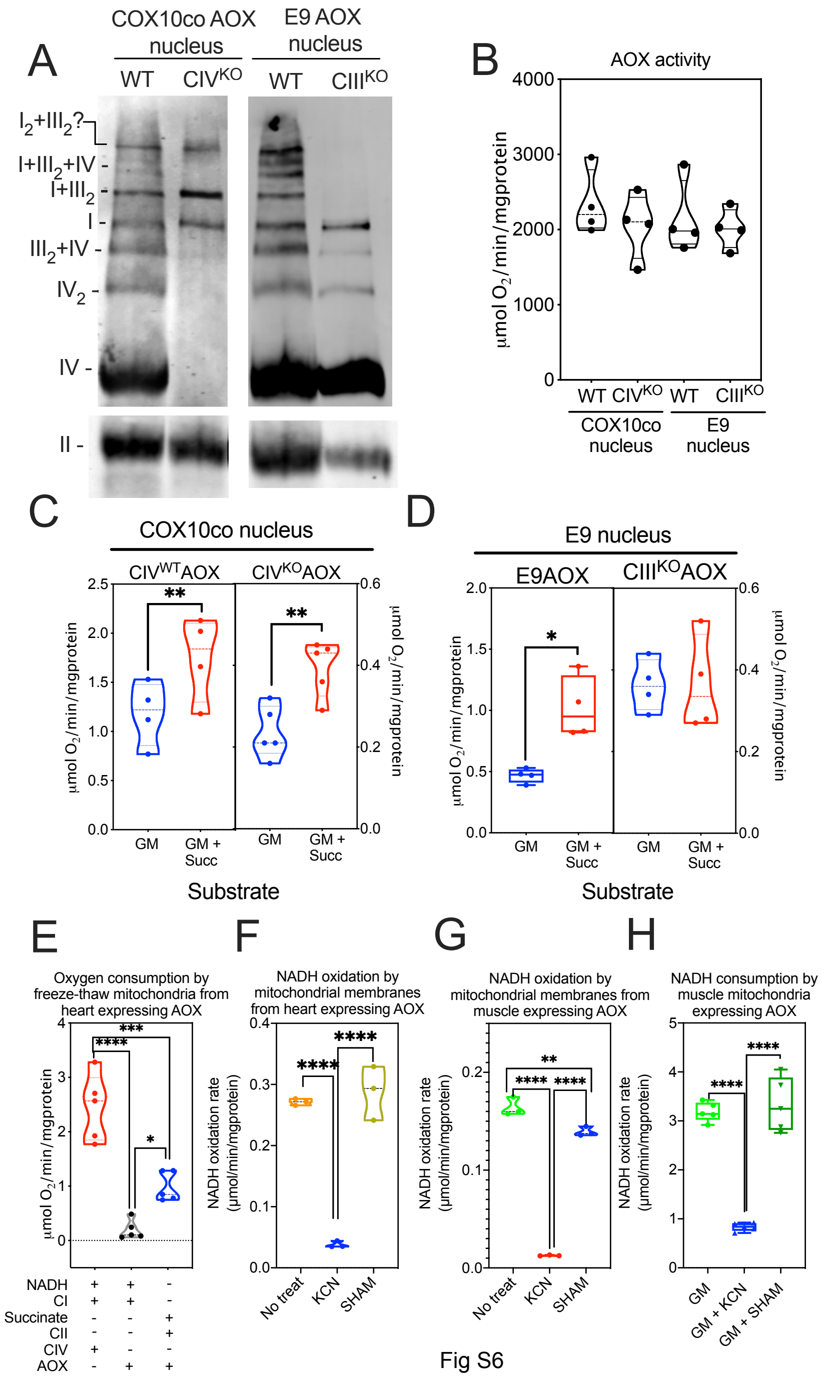


Fig S6

**Figure S6.- AOX as a tool to investigate the segmentation of the CoQ pool. A,** BNGE of mitochondrial preparations of the indicated cell line and proved with antibodies against complexes I, II and IV to illustrate the maintenance of the SC: $I+III_2$ in the absence of CIV and the presence of only free CI in the absence of CIII. **B,** the different cell lines expressing AOX reached a similar level of activity of this enzyme. **C-D**, The increase in respiration by the addition of succinate to mitochondria fed with CI substrates required the formation of SC: $I+III_2$. **E-F**, direct assessment of NADH-dependent respiration (**e**) or NADH oxidization (**f**) by permeabilized heart mitochondria expressing AOX in the indicated conditions. **G-H**, direct assessment of NADH-dependent respiration (**G**) or NADH oxidization (**H**) by permeabilized skeletal muscle mitochondria expressing AOX in the indicated conditions.
